# Robust information routing by dorsal subiculum neurons

**DOI:** 10.1101/2020.06.04.133256

**Authors:** Takuma Kitanishi, Ryoko Umaba, Kenji Mizuseki

## Abstract

The hippocampus conveys various information associated with spatial navigation; yet, information distribution to multiple downstream areas remains unknown. We investigated this by identifying axonal projections using optogenetics during large-scale recordings from the rat subiculum, the major hippocampal output structure. Subicular neurons demonstrated a noise-resistant representation of place, speed, and trajectory, which was as accurate as that of hippocampal CA1 neurons. Speed and trajectory information was most prominently sent to the retrosplenial cortex and nucleus accumbens, respectively. Place information was distributed uniformly to the retrosplenial cortex, nucleus accumbens, anteroventral thalamus, and medial mammillary body. Information transmission by projection neurons was tightly controlled by theta oscillations and sharp-wave/ripples in a target region-specific manner. In conclusion, the subiculum robustly routes diverse navigation-associated information to downstream areas.

**One Sentence Summary:** The subiculum accurately and robustly represents diverse information and routes it in a target region-specific manner.

## Main text

The hippocampus is crucially involved in spatial navigation and memory. Through interaction with intra- and extra-hippocampal areas, the hippocampal neurons acquire diverse representation of spatial navigation parameters, such as place (*1*), speed (*2*), trajectory (*3*), head direction (*4*), and time (*5*). The spike timing of these neurons is secured by multiple hippocampal neural oscillations, including theta oscillations, gamma oscillations, and sharp-wave/ripples (SPW-Rs) (*6–8*), thus facilitating temporally precise information transmission within and between brain regions. To support the navigational behavior output, hippocampal information should be distributed to extrahippocampal areas. However, little is known about how diverse hippocampal information is distributed to downstream areas.

The subiculum (SUB) is the major hippocampal output structure, receiving hippocampal CA1 and entorhinal cortical output and projecting to various cortical/subcortical areas, including the nucleus accumbens (NAC), anteroventral thalamic nucleus (AV), interanteromedial thalamic nucleus (ITN), retrosplenial cortex (RSC), and medial mammillary body (MMB) (*9*). The CA1–SUB projection is topographically organized so that proximal and distal CA1 areas project to the distal (far from CA1) and proximal (close to CA1) SUB, respectively (*9*). The CA1 area shows graded spatial tuning along its proximodistal axis (*10*). In addition, SUB neurons located at proximal and distal areas innervate different target areas (*11*), suggesting that these projections potentially convey distinct information. Consistent with this idea, suppression of the proximal or distal SUB has a distinct impact on memory acquisition (*12*). Analogous to CA1 neurons, SUB neurons represent multiple variables associated with spatial navigation, such as place (*13–15*), axis (*16*), boundary (*17*), reward (*18*), and memory (*19*). Such anatomical and physiological evidence imply a pivotal role for SUB in the distribution of multimodal information; yet, little is known about the nature of information distribution from this structure. An enigmatic point of subicular representation is that the overall subicular spatial firing is apparently dispersed (*13, 14*). Thus, it remains elusive whether the SUB fails to inherit the information content from the CA1 area or whether it inherits the information but convert it into a distinct form of representation suitable for interregional distribution to downstream areas. Moreover, it is unknown whether the SUB distributes multiple types of information uniformly to all target areas or selectively routes distinct information to different targets. We anticipated that the SUB processes navigational information received from the CA1 area and routes the information to downstream targets in a projection-specific manner.

We investigated the above hypothesis by identifying axonal projections using optogenetics (*20*) while performing large-scale extracellular recordings (*21*) of rat dorsal SUB and CA1 area during multiple spatial tasks and sleep. We found that the SUB has a noise-resistant, accurate representation of multiple types of navigation-associated information and distributes this information uniformly or selectively to target areas depending on the type of information.

## Results

### Anatomical tracing

Anterograde tracing revealed that the major projection targets of the rat dorsal SUB include NAC, AV, anterodorsal thalamic nucleus, ITN, nucleus reuniens, RSC, and MMB (Fig. S1). Moderate projection was also observed in the dorsal tenia tecta, dorsal peduncular cortex, and entorhinal cortex (Fig. S1). These projections were consistent with those reported by previous studies (*11*). Next, we determined the somata locations of SUB projection neurons by injecting the retrograde tracer cholera toxin B subunit conjugated with Alexa Fluor 488 (CTB488) into one of the above target areas. In addition to the well-characterized differences between the proximal and distal SUB areas (*11*), we observed robust superficial-deep differences in the locations of SUB projection neurons (Fig. 1A, Fig. S2, S3). Projection-specific visualization of neuronal morphology (*22*) revealed that dendritic spine density was lower in AV-projecting (AV-p) and ITN-projecting (ITN-p) neurons than in NAC-projecting (NAC-p) and RSC-projecting (RSC-p) neurons (Fig. S4). These results, together with accumulating previous evidence (*11, 12*), indicate that SUB projection neurons comprise at least partly-separated groups of neurons that target distinct downstream areas.

**Fig. 1.**
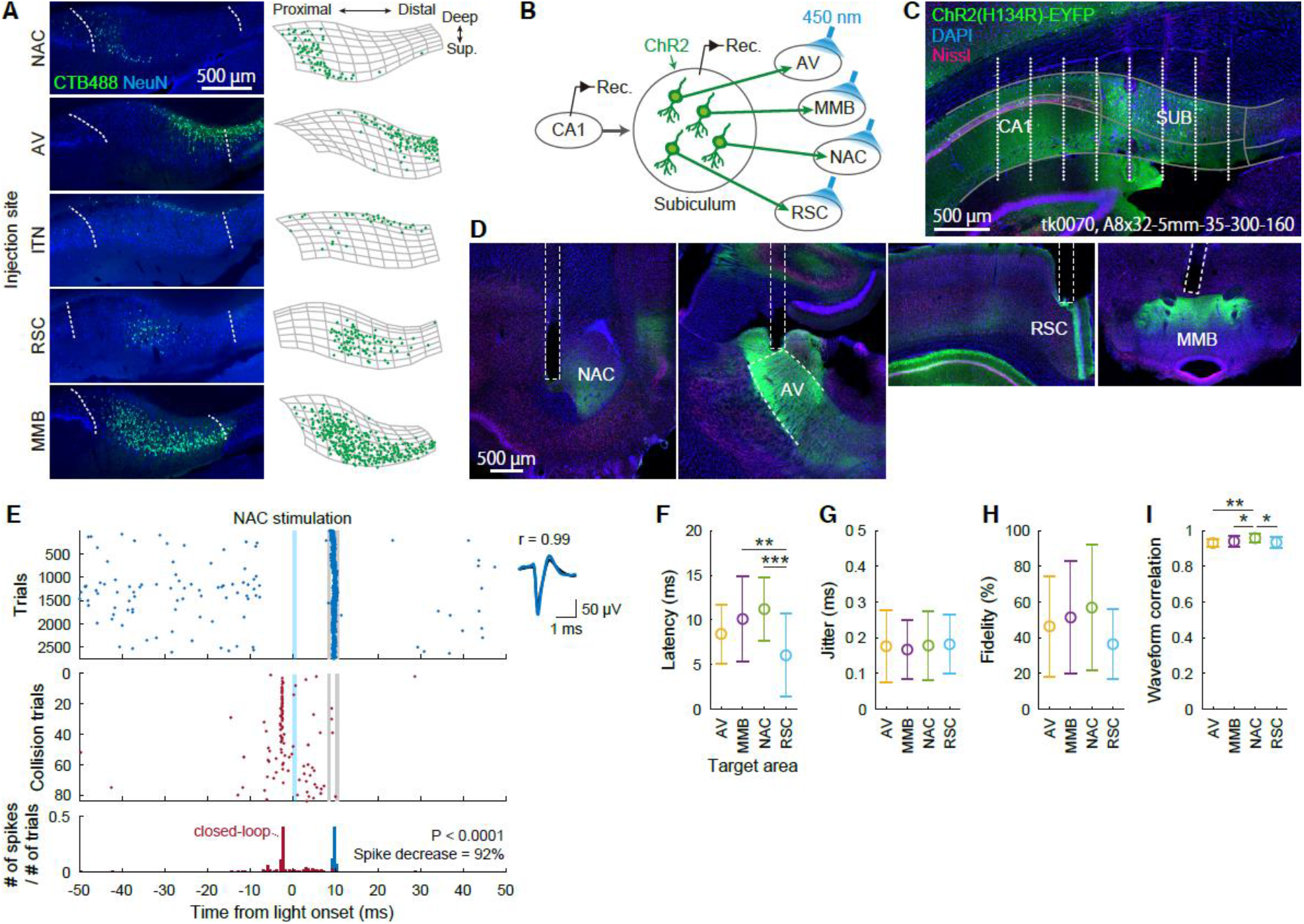
Optogenetic identification of SUB projection neurons. (A) SUB coronal sections containing CTB488-labeled projection neurons (left) and the corresponding cell counting (right). (Left) Dotted curves: borders of SUB cell layer. (Right) Segmented SUB cell layer (mesh) and detected CTB488-positive neurons (dots). (B) Schematic of optogenetic identification. (C) Coronal section indicating ChR2 expression (green) and the estimated recording sites (white dots). Gray curves: borders of CA1, SUB, and their cell layers. Text: animal ID and silicon probe name. (D) NAC, AV, RSC, and MMB coronal sections (from left to right) containing ChR2-expressing SUB axons (green) and the reconstructed optical fiber locations (dotted lines). (E) Left: Spike responses of a SUB neuron upon light irradiation to the NAC. Evoked spikes (top) decreased when spontaneous spikes occurred shortly before the expected latency of evoked spikes (middle, bottom). Shaded blue areas: light stimulation periods. Gray lines: bin ranges of evoked spikes. Right: Mean waveforms of spontaneous (black) and evoked (blue) spikes. (F–H) Latency (F), jitter (G), and fidelity (H) of evoked spikes. (I) Waveform correlation between the evoked and spontaneous spikes. (F–I) Mean ± SD. * *P* < 0.05, ** *P* < 0.01, *** *P* < 0.001, Tukey test.

### Optogenetic identification of projection targets

The anatomical observations led us to hypothesize that SUB projection neurons distribute distinct information to different projection targets. To investigate this, we combined large-scale extracellular recordings (*21*) from the SUB and CA1 area of freely-behaving rats and optogenetic identification of the projection targets (*20*) of the recorded SUB neurons. We stereotaxically introduced adeno-associated virus (AAV) expressing channelrhodopsin-2 under the control of synapsin promoter, AAV1-hSyn-hChR2(H134R)-EYFP, into the dorsal SUB, inserted a 256-channel silicon probe into the dorsal SUB and CA1 area, and implanted up to four optical fibers to each projection target (NAC, AV, RSC, and MMB) in each rat (Fig. 1B–1D, Fig. S5). Rats performed multiple spatial tasks, including an open-field task, a linear-track task, and an alternating T-maze task, and rested/slept in a small enclosure before and after the tasks. We monitored and analyzed a total of 315 and 319 putative principal cells in the CA1 and SUB, respectively, from 22 recording sessions in 11 rats (Fig. S6).

To identify the projection targets of activity-monitored neurons with antidromic spikes (*20*), we irradiated blue light pulses sequentially to each target area during the rest sessions. In response to the light pulses, a proportion of SUB neurons generated short-latency (<25 ms), low-jitter (<0.5 ms) spikes with high fidelity (>20%) and were identified as projection neurons innervating the irradiated area (Fig. 1E–1I, Fig. S7). The locations of the identified projection neurons in the SUB cell layer were consistent with the distribution of retrogradely-labeled CTB488-positive neurons (Fig. S7). RSC-p neurons had a shorter latency than NAC-p and MMB-projecting (MMB-p) neurons (Fig. 1F), presumably reflecting differences in the anatomical distance between the SUB and these target areas. Among SUB principal cells, we identified 11 AV-p (3.4% of SUB principal cells), 28 MMB-p (8.8%), 18 NAC-p (5.6%), and 16 RSC-p (5.0%) SUB neurons.

### The SUB represents place information as accurately as the CA1 and distributes this information uniformly

The hippocampal CA1 area and the SUB contain spatially tuned cells that fire whenever animals move through particular places in an environment (*1, 13–15*). We compared place representation in CA1 and SUB neurons while rats were running on a linear track. While many CA1 neurons fired at specific places along the track, SUB neurons fired at broader locations (Fig. 2A). Consistent with our visual inspection results, per-spike spatial information, expressed by the *I_spike_* value, a measure of the spatial specificity of individual cells (*23*), was lower in SUB than in CA1 neurons [mean ± standard deviation (SD): CA1, 0.94 ± 0.77 bits/spike; SUB, 0.34 ± 0.46 bits/spike; *Z* = 9.73, *P* < 0.0001, Wilcoxon rank sum test; Fig. 2B]. In contrast, SUB neurons had higher mean firing rates (during trials in the preferred direction; CA1, 3.7 ± 5.4 Hz; SUB, 9.8 ± 12.9 Hz; *Z* = 7.78, *P* < 0.0001; Fig. 2C) and peak firing rates (in the preferred direction; CA1, 9.3 ± 9.0 Hz; SUB, 17.1 ± 17.0 Hz; *Z* = 5.83, *P* < 0.0001; Fig. 2D) than CA1 neurons. Due to this high firing rate, per-second spatial information, *I_sec_* (*23*), which is equal to *I_spike_* multiplied by the mean firing rate, was comparable between CA1 and SUB neurons (CA1, 1.43 ± 1.69 bits/s; SUB, 1.39 ± 1.87 bits/s; *Z* = 0.98, *P* = 0.33; Fig. 2E). Moreover, SUB and CA1 neurons showed similar mutual information (*24*) of firing rates and places (CA1, 0.26 ± 0.17 bits; SUB, 0.29 ± 0.23 bits; *Z* = 0.51, *P* = 0.61; Fig. 2F). The mutual information correlated with *I_sec_* (CA1: *r* = 0.87, *P* < 0.0001; SUB: *r* = 0.84, *P* < 0.0001) but not with *I_spike_* (CA1: *r* = 0.004, *P* = 0.96; SUB: *r* = 0.11, *P* = 0.08). These results suggest that, despite the apparent decrease in spatial specificity as shown by the lower *I_spike_* value, SUB neurons convey a comparable amount of spatial information per unit time with that conveyed by CA1 neurons owing to the subicular high firing rate. Theoretical modeling suggested that the CA1 and SUB employ different strategies to maximize *I_sec_* (Fig S8). The idea that SUB neurons robustly represent place information is further supported by the observation that their firing was anchored more to place than to the elapsed time from the start of trials (Fig. S9A–D), and by the fact that SUB rate maps were more temporally stable than the CA1 rate maps (Fig. S9E).

**Fig. 2.**
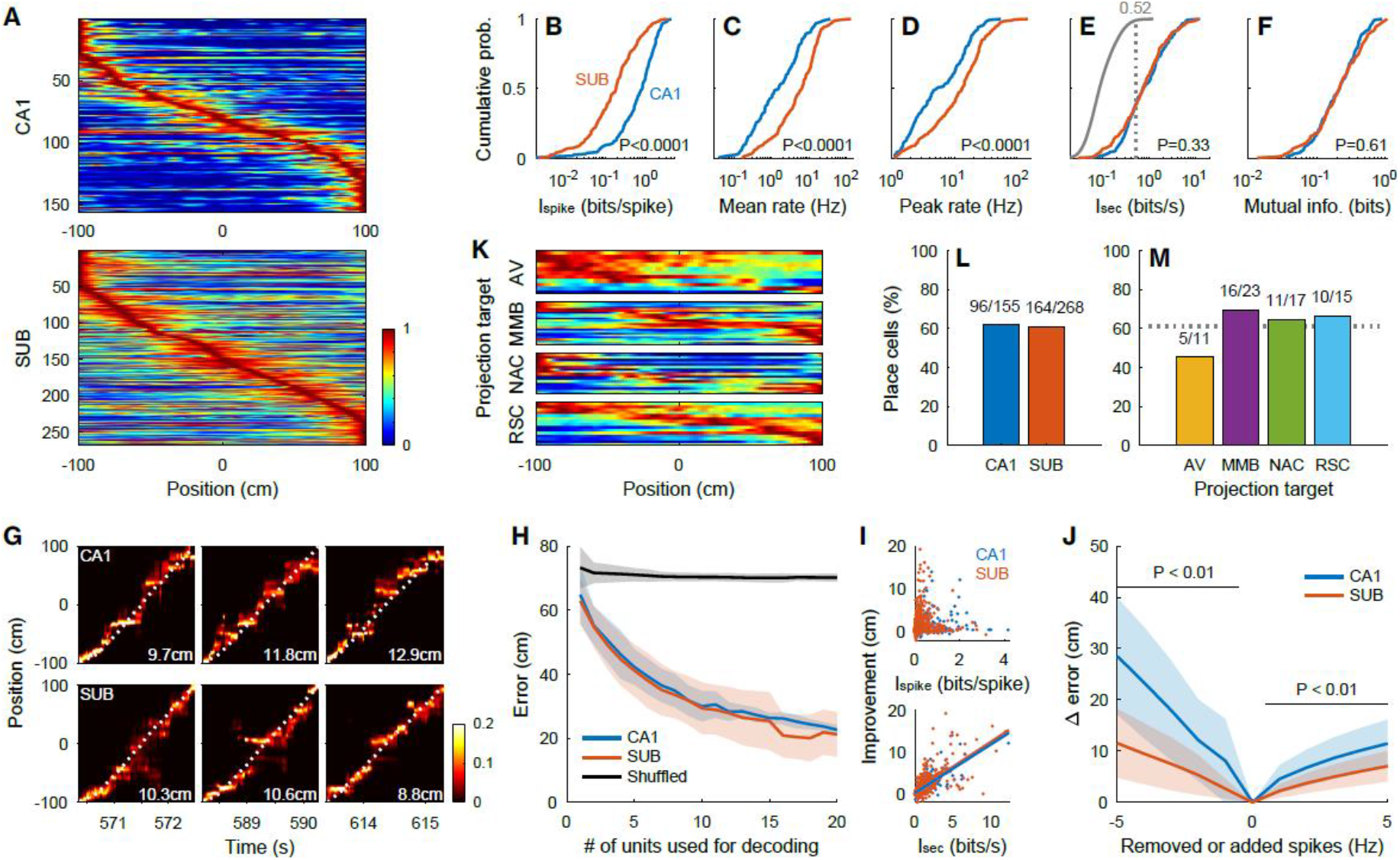
Place representation on a linear track. (A) Peak-normalized rate maps of CA1 (top) and SUB (bottom) neurons sorted by peak positions. (B–F) Cumulative distribution of *I_spike_* (B), mean firing rate (C), peak firing rate (D), *I_sec_* (E), and mutual information (F) of CA1 and SUB neurons. *P* values: Wilcoxon rank sum test. (E) The gray curve, number, and dotted line: the distribution of shuffled data and its 99th percentile. (G) Position decoding from simultaneously recorded CA1 (top) or SUB (bottom) neurons in three consecutive trials. Color maps: probabilities of estimated animal positions. Dotted curves: observed animal positions. Numbers: decoding errors of the trials. (H) Error of position decoding. Black line: decoding error obtained from a shuffling procedure. (H, J) Mean (solid lines) ± SD (shaded areas). (I) Decoding improvement plotted against *I_spike_* (top) and *I_sec_* (bottom). Lines: linear regression. (J) Difference in decoding errors after randomly removing (negative x values) or adding (positive x values) spikes to each neuron. *P* < 0.01, Bonferroni test after two-way repeated-measures ANOVA. (K) Peak-normalized rate maps of SUB projection neurons sorted by peak positions. (L, M) Percentage of place cells in the CA1 and SUB (L), and SUB projection neurons (M). Numbers: number of place cells to the total number of neurons. Dotted line: percentage of place cells in SUB neurons.

Next, we investigated whether the SUB also represents place information as accurately as the CA1 at a cell population level using Bayesian decoding of animal position (*25*) (Fig. 2G–H). The decoding errors obtained from CA1 and SUB neuronal populations were both smaller than those obtained using shuffled data, even when single neurons were used for decoding (*P* < 0.001, Tukey test), and became even smaller by increasing the numbers of neurons used [*F*19, 700 = 90.87, *P* < 0.001, two-way analysis of variance (ANOVA); Fig. 2H]. Importantly, the decoding errors were comparable between the CA1 and SUB, irrespective of the number of neurons used (*P* > 0.2, Bonferroni test after two-way ANOVA; Fig. 2H), suggesting that CA1 and SUB neuronal populations of the same size convey a comparable amount of decodable spatial information.

To clarify how the spatial firing of single neurons relates to population coding, we estimated the contribution of single neurons to the decoding performance. The decoding improvement by adding single units correlated with *I_sec_* (CA1: *r* = 0.76, *P* < 0.0001; SUB: *r* = 0.69, *P* < 0.0001) and mutual information (CA1, *r* = 0.78, *P* < 0.0001; SUB, *r* = 0.73, *P* < 0.0001), but not with *I_spike_* (CA1, *r* = 0.06, *P* = 0.46; SUB, *r* = 0.04, *P* = 0.40) in both CA1 and SUB (Fig. 2I). In addition, increasing *I_sec_* by increasing the firing rate but without changing the spatial tuning improved the decoding performance (Fig. S10), highlighting the tight association between *I_sec_* and population coding.

We hypothesized that the subicular representation gains robustness against noises owing to its high firing rate. We tested this idea by adding noises to the original spike trains before proceeding with the decoding analysis. In response to simple additive noises (*26*), we found a smaller increase in the decoding error in the SUB than in the CA1 (Fig. 2J), suggesting that population spatial coding in the SUB is more resistant to additive noises than that in the CA1.

Next, we asked how spatial information in the SUB is distributed to its downstream targets (Figs. 2K–M). The proportions of place cells (defined by *I_sec_*) were similar in the CA1 and SUB (*χ*^2^ = 0.02, *P* = 0.88, *χ*^2^ test; Fig. 2L), as well as among the groups of SUB projection neurons targeting the AV, MMB, NAC, or RSC (*χ*^2^ = 1.99, *P* = 0.57; Fig. 2M), suggesting that spatial information is distributed uniformly from the SUB to each of these four downstream targets. We observed a similar spatial representation in a two-dimensional open field (Fig. S11).

### SUB represents speed information more accurately than CA1 and send the information to RSC

The hippocampus contains neurons that respond to the running speed of animals (*2*). We thus investigated speed representation in CA1 and SUB neurons while rats foraged in an open field. Both the CA1 and SUB contained neurons with instantaneous firing rates that correlated with the rat’s instantaneous running speed (Fig. 3A). To estimate the amount of speed information, as in the case of spatial information, we calculated per-spike speed information *I_spike_* and per-second speed information *I_sec_* (*27*). The *I_spike_* value was lower in SUB than in CA1 neurons (CA1, 0.054 ± 0.057 bits/spike; SUB, 0.028 ± 0.059 bits/spike; *Z* = 9.12, *P* < 0.0001), while the mean firing rates of SUB neurons were higher than those of CA1 neurons (CA1, 1.5 ± 1.9 Hz; SUB, 5.6 ± 5.2 Hz; *Z* = 11.48; *P* < 0.0001) (Fig. 3B–C). Consequently, the *I_sec_* was similar between all CA1 and SUB neurons (CA1, 0.046 ± 0.052 bits/s; SUB, 0.077 ± 0.114 bits/s; *Z* = 1.12; *P* = 0.26; Fig. 3D). However, when comparing only neurons with an *I_sec_* exceeding the 99th percentile of the shuffled data, the *I_sec_* was higher for SUB than for CA1 neurons (CA1, 0.118 ± 0.060 bits/s; SUB, 0.205 ± 0.137 bits/s; *Z* = 5.19, *P* < 0.0001; Fig. 3D). To distinguish positively- and negatively-modulated speed cells, we calculated the speed score (*28*). The absolute speed score tightly correlated with *I_sec_* (CA1, *r* = 0.75, *P* < 0.0001; SUB, *r* = 0.80, *P* < 0.0001) but only modestly correlated with *I_spike_* (CA1, *r* = 0.32, *P* < 0.0001; SUB, *r* = 0.32, *P* < 0.0001). The speed score was higher for SUB than for CA1 neurons (CA1, 0.141 ± 0.043; SUB, 0.201 ± 0.087; *Z* = 5.74, *P* < 0.0001) when comparing neurons with a speed score exceeding the 99th percentile of the shuffled data distribution (Fig. 3E). Conversely, the speed score was smaller for SUB than for CA1 neurons (CA1, –0.100 ± 0.006; SUB, –0.142 ± 0.041; *Z* = 2.34, *P* = 0.019) when comparing neurons with a speed score smaller than the 1st percentile of the shuffled data distribution (Fig. 3E). The speed scores obtained from the first and second halves of the sessions were positively correlated both for the CA1 and the SUB (CA1, *r* = 0.51, *P* < 0.0001; SUB, *r* = 0.79, *P* < 0.0001). These results suggest that, despite the broader speed tuning, as shown by the lower *I_spike_* values, SUB neurons convey a larger amount of speed information per unit time than that conveyed by CA1 neurons.

**Fig. 3.**
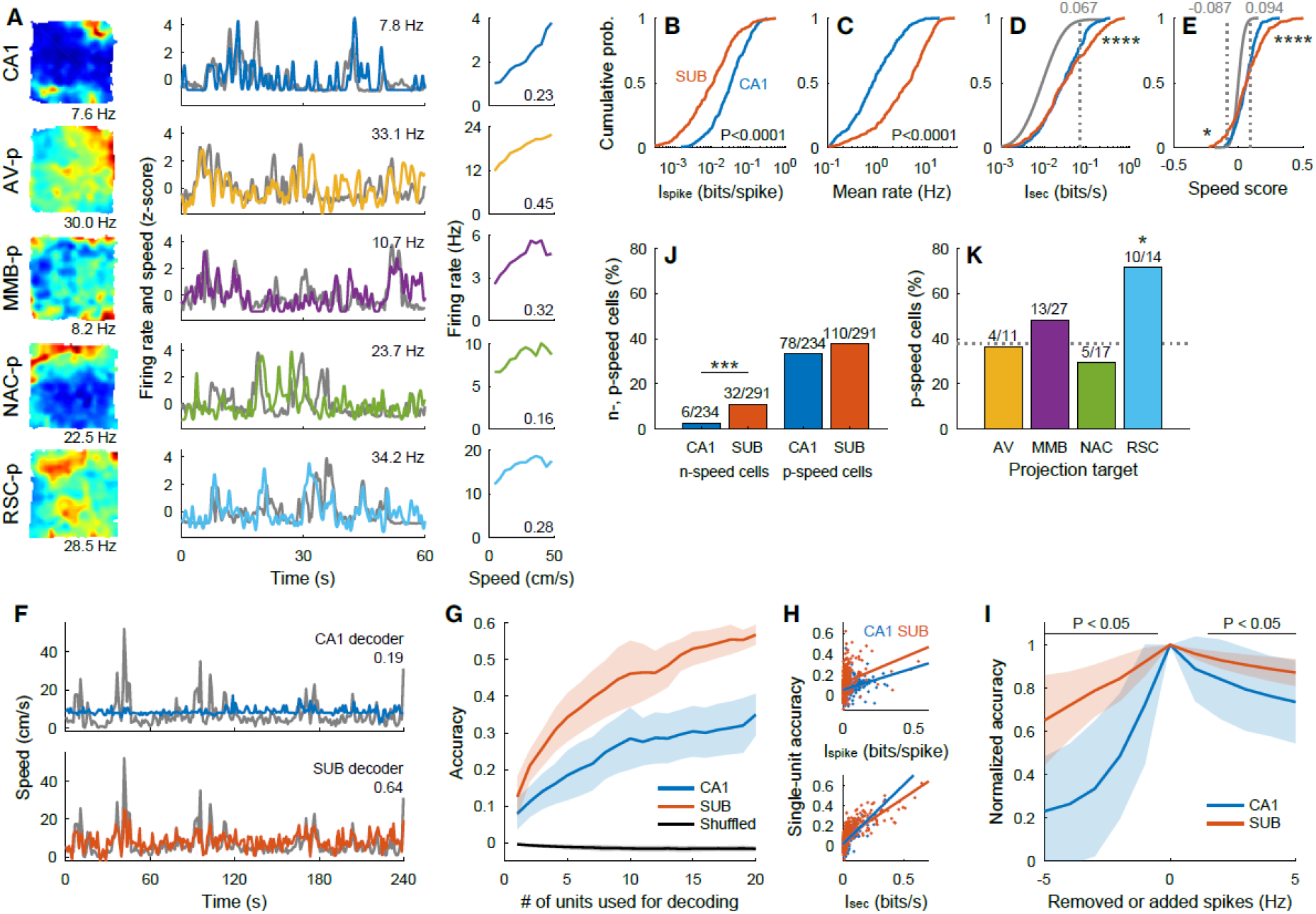
Speed representation in an open field. (A) Rate maps (left), z-scored running speed (gray) and firing rate (color) (middle), and mean firing rate as a function of running speed (right). Each row shows the firing patterns of a single neuron. (left, middle) Numbers: peak firing rates in the figure. (right) Numbers: speed scores. (B–E) Cumulative distribution of *I_spike_* (B), mean firing rates (C), *I_sec_* (D), and speed scores (E). *P* values: Wilcoxon rank sum test. (D, E) Gray curves, numbers, and dotted lines: the distribution of the shuffled data and its 1st (E) and 99th (D, E) percentiles. * *P* = 0.019, **** *P* < 0.0001, CA1 *vs*. SUB neurons exceeding either the 1st- or 99th-percentile threshold, Wilcoxon rank sum test. (F) Decoding of speed from simultaneously recorded CA1 (top) and SUB (bottom) activity. Gray: observed speed. Color: decoded speed. Numbers: decoding accuracy. (G) Decoding accuracy. Black line: decoding error obtained from a shuffling procedure. (G, I) Mean (solid lines) ± SD (shaded areas). (H) Single-unit decoding accuracy plotted against *I_spike_* (top) and *I_sec_* (bottom). Lines: linear regression. (I) Normalized decoding accuracy after randomly removing (negative x values) or adding (positive x values) spikes to each neuron. *P* < 0.05, Bonferroni test after two-way repeated-measures ANOVA. (J, K) Percentage of n-speed (left) and p-speed (right) cells in the CA1 and SUB (J), and percentage of p-speed cells in SUB projection neurons (K). Numbers: number of speed cells to the total number of neurons. (J) *** *P* = 0.0002, χ^2^ test. (K) Dotted line: percentage of p-speed cells in SUB neurons. \**P* < 0.05, bootstrap analysis.

We asked whether the stronger speed representation in individual SUB neurons is reflected by the speed representation at the population level. To this end, we constructed a linear decoder of speed, as described previously (*28*) (Fig. 3F–G). The decoding accuracy was higher in the SUB than the CA1 irrespective of the number of neurons used (*P* < 0.05, Bonferroni test after two-way ANOVA; Fig. 3G). The single-unit decoding accuracy better correlated with the *I_sec_* (CA1: *r* = 0.68, *P* < 0.0001; SUB: *r* = 0.75, *P* < 0.0001) than with *I_spike_* (CA1: *r* = 0.27, *P* < 0.0001; SUB: *r* = 0.28, *P* < 0.0001) both in the CA1 and SUB (CA1: *Z* = 5.85, *P* < 0.0001; SUB: *Z* = 8.03, *P* < 0.0001; test of the difference between two correlation coefficients; Fig. 3H). Speed decoding by the SUB was more resistant to additive noises than that by the CA1 (Fig. 3I).

We then investigated the proportion of positively- (p-speed cells) and negatively- (n-speed cells) modulated speed cells (see Materials and Methods). While the proportion of p-speed cells was similar between the SUB and CA1 (*χ*^2^ = 1.12, *P* = 0.29; Fig. 3J), that of n-speed cells was higher in the SUB than in the CA1 (*χ*^2^ = 13.71, *P* = 0.0002; Fig. 3J). RSC-p neurons contained a larger proportion of p-speed cells than that of the entire SUB cell population (*P* < 0.05, bootstrap analysis; Fig. 3K). This result indicates that speed information in the SUB is most prominently routed to the RSC.

### SUB represents trajectory information as accurately as the CA1 and send this information to NAC

Next, we investigated trajectory-dependent firing at the start point of alternating T-maze task (Fig. 4A) (*3*). A fraction of CA1 and SUB neurons showed different firing rates in the start box according to the next choice of arm (Fig. 4B). To measure the information encoded about the next choice (left or right), signaled by the firing rate in the start box, we defined trajectory *I_spike_* and *I_sec_*. Compared with CA1 neurons, SUB neurons showed lower *I_spike_* values (CA1, 0.034 ± 0.064 bits/spike; SUB, 0.011 ± 0.023 bits/spike; *Z* = 5.73, *P* < 0.0001; Fig. 4C) and lower rate change ratio (CA1, 0.24 ± 0.19; SUB, 0.15 ± 0.13; *Z* = 5.72, *P* < 0.0001; Fig. 4D). However, SUB neurons had higher mean firing rate in the start box (CA1, 2.3 ± 3.2 Hz; SUB, 6.8 ± 6.8 Hz; *Z* = 10.16, *P* < 0.0001; Fig. 4E), and consequently had comparable *I_sec_* values to those of CA1 neurons (CA1, 0.034 ± 0.063 bits/s; SUB, 0.040 ± 0.089 bits/s; *Z* = 0.02, *P* = 0.99; Fig. 4F). In addition, the area under the receiver operating characteristic (auROC), indicating the goodness of binary classifier, was similar between CA1 and SUB neurons (CA1, 0.601 ± 0.084; SUB, 0.609 ± 0.091; *Z* = 0.92, *P* = 0.36; Fig. 4G). We performed decoding of trajectory from the firing rates in the start box, as described previously (*29*). SUB neurons showed moderately higher decoding accuracy than that of CA1 neurons (*F*_1, 432_ = 17.02, *P* < 0.001; Fig. 4H) and were more resistant to additive noise (Fig. 4I).

**Fig. 4.**
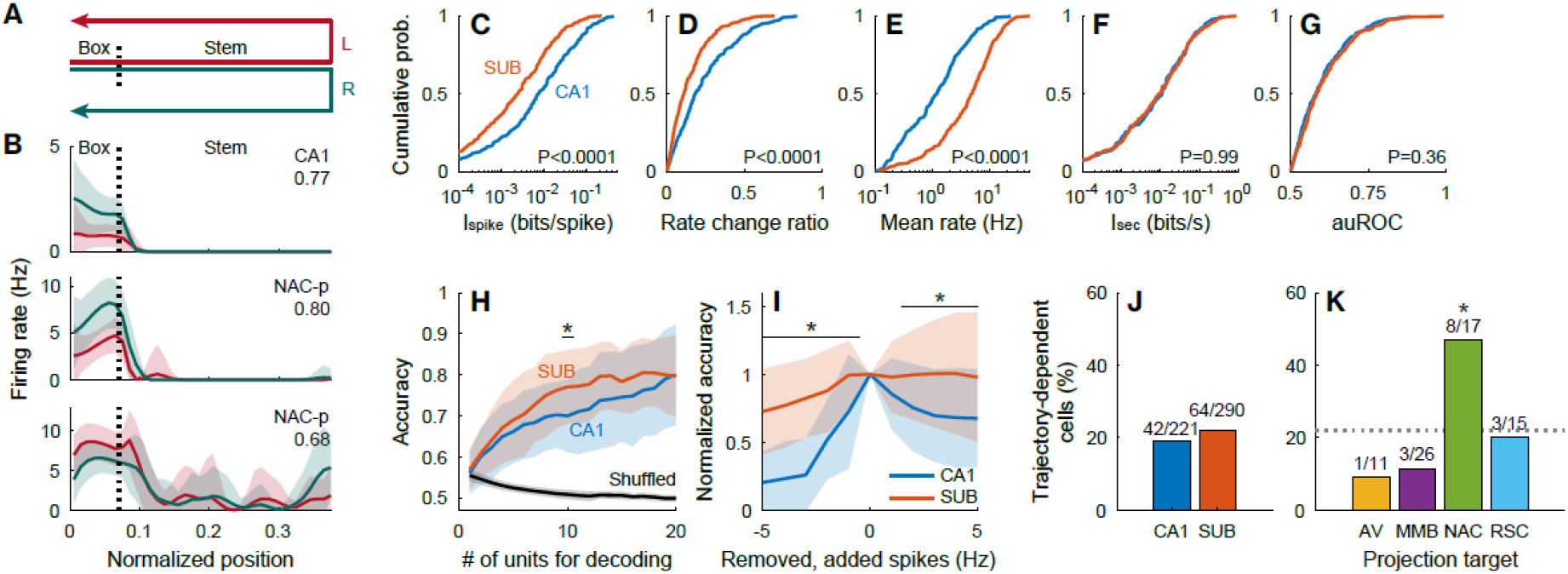
Trajectory-dependent firing during an alternating T-maze task. (A) Schematic of the behavioral task. (B) Rate maps of a CA1 and two NAC-p SUB neurons while rats were in the start box (normalized position, 0.02–0.07) and stem (normalized position, 0.07–0.38). Numbers: auROC of the neuron. Red: left-arm trials. Green: right-arm trials. Mean (solid lines) ± SD (shaded areas). (C–G) Cumulative distribution of *I_spike_* (C), mean rate change (D), mean firing rate (E), *I_sec_* (F), and auROC (G) in the start box. *P* values: Wilcoxon rank sum test. (H) Accuracy of trajectory decoding. Black line: chance level of decoding error estimated by a shuffling procedure. * *P* < 0.05, Bonferroni test after two-way ANOVA. (H, I) Mean (solid lines) ± SD (shaded areas). (I) Normalized decoding accuracy after randomly removing (negative x values) or adding (positive x values) spikes to each neuron. * *P* < 0.05, Bonferroni test after two-way repeated-measures ANOVA. (J, K) Percentage of trajectory-dependent cells in the CA1 and SUB (J), and SUB projection neurons (K). Numbers: number of trajectory-dependent cells to the total number of neurons. Dotted line: percentage of speed cells in SUB neurons. * *P* < 0.05, bootstrap analysis.

The CA1 and SUB contained similar proportions of trajectory-dependent cells (*χ*^2^ = 0.72, *P* = 0.40; Fig. 4J). Among groups of SUB projection neurons, NAC-p neurons contained a higher proportion of trajectory-dependent cells than the other groups of projection neurons (*χ*^2^ = 9.06, *P* = 0.03) and the entire population of SUB neurons (*P* < 0.05, bootstrap analysis; Fig. 4K).

### Projection-specific phase locking to theta oscillations

Next, we asked whether the CA1, SUB, and SUB projection neurons have specific temporal firing patterns during theta oscillations. During running, we observed robust theta oscillations both in the CA1 and SUB (Figs. 5A–5C). Using theta oscillations at the center of the SUB cell layer as a reference, we examined theta phase locking of neuronal firing during RUN periods. Virtually all CA1 and SUB neurons were significantly phase-locked (*P* < 0.05, Rayleigh test) to subicular theta oscillations (CA1, 100%; SUB, 99.0%; Fig. 5D). Preferred theta phases of SUB neurons were at earlier phases of theta cycles than those of CA1 neurons (CA1, 46.1 ± 45.4 degrees; SUB, 332.6 ± 54.7 degrees; circular mean ± angular deviation; *U*^2^ = 4.30, *P* < 0.001, Watson *U*^2^ test; Fig. 5D–5E). We measured the strength of phase locking using pairwise phase consistency (PPC) (*30*). While CA1 neurons locked to the ascending phase of theta oscillations showed higher PPC than those locked to the descending phase (*F*_9,267_ = 4.80, *P* < 0.0001, one-way ANOVA), SUB neurons locked to the peaks of theta oscillations showed higher PPC than those locked to troughs (*F*_9, 290_ = 2.69, *P* = 0.005) (Fig. 5D). As the entire population, PPC was modestly smaller in the SUB than in the CA1 (CA1, 0.060 ± 0.058; SUB, 0.052 ± 0.065; *Z* = 2.94; *P* = 0.003).

**Fig. 5.**
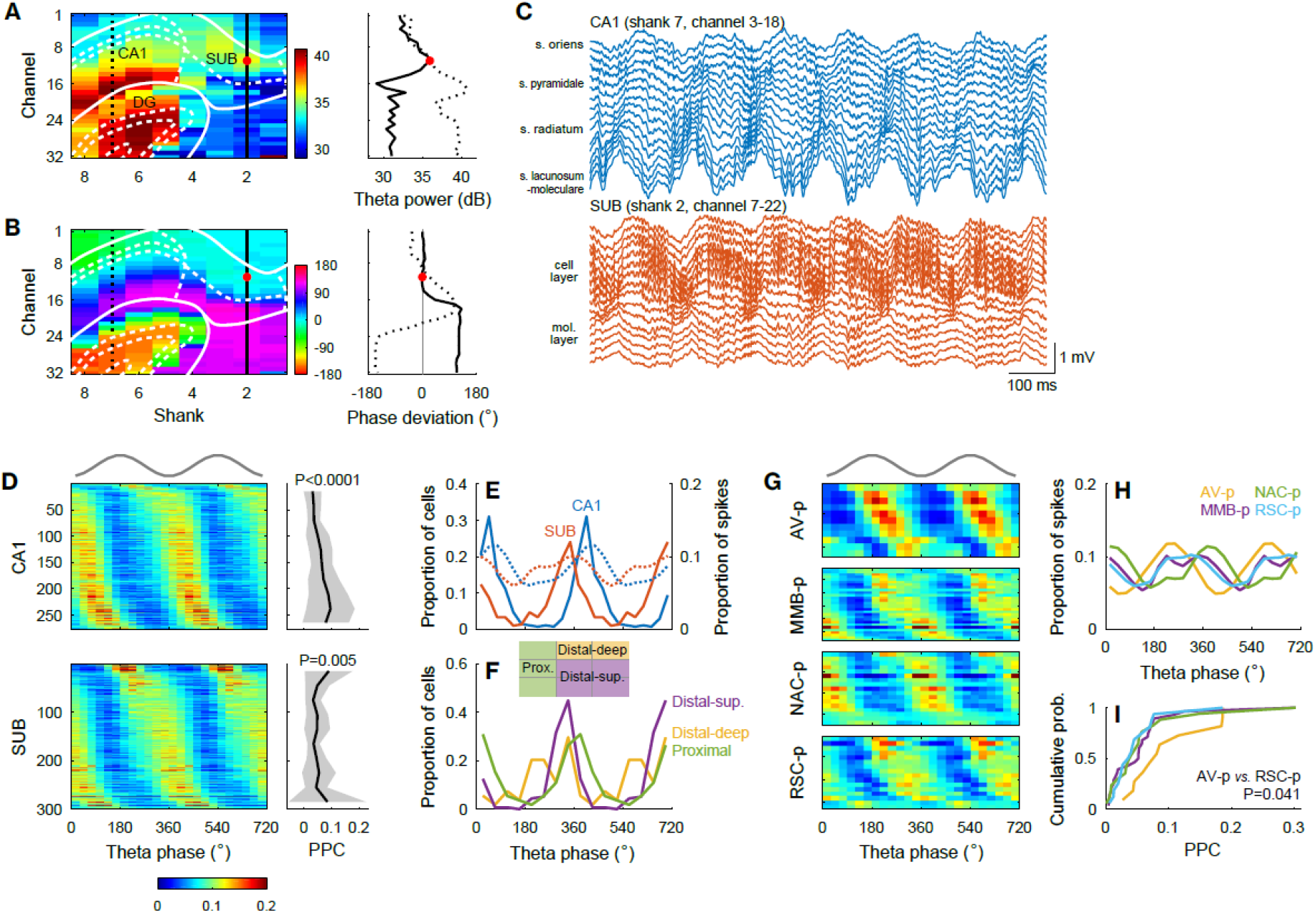
Projection-specific theta phase locking during RUN periods. (A, B) Distribution of theta power (A) and theta phase deviations (B) in an open field (left). White lines: borders of the CA1, SUB, and DG. Values along the CA1 (dotted black lines) and SUB (solid black lines) are plotted in the right panels. Red dots: the reference recording site with the maximal theta power in the SUB cell layer. (C) Wide-band local field potentials simultaneously recorded from the CA1 (top) and SUB (bottom). (D) Spike phase distribution along the reference SUB theta oscillations (left) and corresponding PPC (right) in the CA1 (top) and SUB (bottom). P values, one-way ANOVA. Mean (solid lines) ± SD (shaded areas). (D, G) Top gray traces: idealized reference theta cycles. (E) Distribution of the preferred theta phases (left y axis, solid lines) and spike theta phases (right y axis, dotted lines). (F) Distribution of preferred theta phases for neurons located at proximal (green), distal-deep (yellow), and distal-superficial (purple) part of the SUB cell layer. (G) Spike phase distribution along the reference SUB theta oscillations in SUB projection neurons. (H, I) Distribution of the spike theta phases (H) and PPC (I) of SUB projection neurons.

Interestingly, SUB neurons had distinct preferred theta phases depending on the location of their somata. SUB neurons located at the distal-deep part (distal two-thirds and deep one-third of the cell layer) of the SUB cell layer fired at the earliest phases of theta cycles (285.4 ± 63.2 degrees), followed by the neurons at distal-superficial part (distal two-thirds and superficial two-thirds of the cell layer, 321.2 ± 39.6 degrees), and finally the neurons at proximal part (proximal one-third of the cell layer, 9.6 ± 54.8 degrees, Fig. 5F). These three distributions of preferred theta phases were different with each other (*P* < 0.01, Watson *U*^2^ test with Bonferroni correction). In agreement with this location-dependent sequential firing, SUB projection neurons showed differences in theta phase locking (Fig. 5G–H). The earliest preferred theta phases were observed in AV-p neurons (250.8 ± 50.0 degrees) followed by RSC-p neurons (292.1 ± 53.6 degrees), MMB-p neurons (297.8 ± 57.0 degrees), and finally NAC-p neurons (29.3 ± 48.6 degrees). The preferred phases of NAC-p neurons were different from those of MMB-p, RSC-p, and AV-p neurons (*P* < 0.02, Watson *U*^2^ test with Bonferroni correction). AV-p neurons were more strongly phase-locked to theta oscillations than RSC-p neurons (*t* = 2.64, *P* = 0.041, Steel-Dwass test; Fig. 5I). These results demonstrate that theta oscillations robustly control the spike timing of SUB neurons in a target-specific manner. We observed similar patterns of theta phase locking during rapid-eye-movement sleep in both the CA1 and SUB (Fig. S12).

### Projection-specific firing modulation by SPW-Rs

Next, we investigated the temporal firing patterns along SPW-Rs during slow-wave sleep (SWS). We observed synchronous SPW-Rs in the CA1 and SUB areas (Fig. 6A–B). Using the ripple events detected at the center of the SUB cell layer as a reference, we examined the peri-ripple event firing rates of individual neurons. While most CA1 (92.7%) and SUB (73.7%) neurons showed significantly (*P* < 0.01, *t*-test) higher firing rates during ripple events than during baseline periods, a small fraction of SUB neurons, but not of CA1 neurons, were significantly (*P* < 0.01, *t*-test) suppressed during ripples (CA1, 0%; SUB, 4.4%; *χ*^2^ = 14.11, *P* = 0.0002; Fig. 6C–D) (*31*). These suppressed neurons were mostly localized in the distal-deep part of the SUB cell layer (Fig. S13). Consistent with this location-dependent firing modulation, 73% of AV-p neurons were significantly suppressed during ripples, while only 18% of them were activated (Fig. 6E). In contrast, nearly all MMB-p (93%) and NAC-p (100%) neurons were activated during ripples (Fig. 6E). RSC-p neurons comprised a mixture of activated (63%) and suppressed (31%) neurons (Fig. 6E). Consequently, AV-p neurons showed the lowest normalized peak height among all projection neuron groups, while RSC-p neurons showed a lower normalized peak height than did MMB-p neurons (Fig. 6F). The ripple-triggered activation / suppression was maintained regardless of the variations in the ripple power (*P* < 0.01 by Bonferroni test after two-way repeated-measures ANOVA, AV-p *vs*. all other projection neuron groups, MMB-p *vs*. RSC-p; *P* = 0.017, NAC-p *vs*. RSC-p; Fig. 6G) or duration (*P* < 0.01, AV-p *vs*. all other projection neuron groups, MMB-p *vs*. RSC-p; *P* = 0.024, NAC-p *vs*. RSC-p; Fig. 6H). Essentially the same pattern of firing modulation was observed in SPW-Rs during the awake rest periods (Fig. S14).

**Fig. 6.**
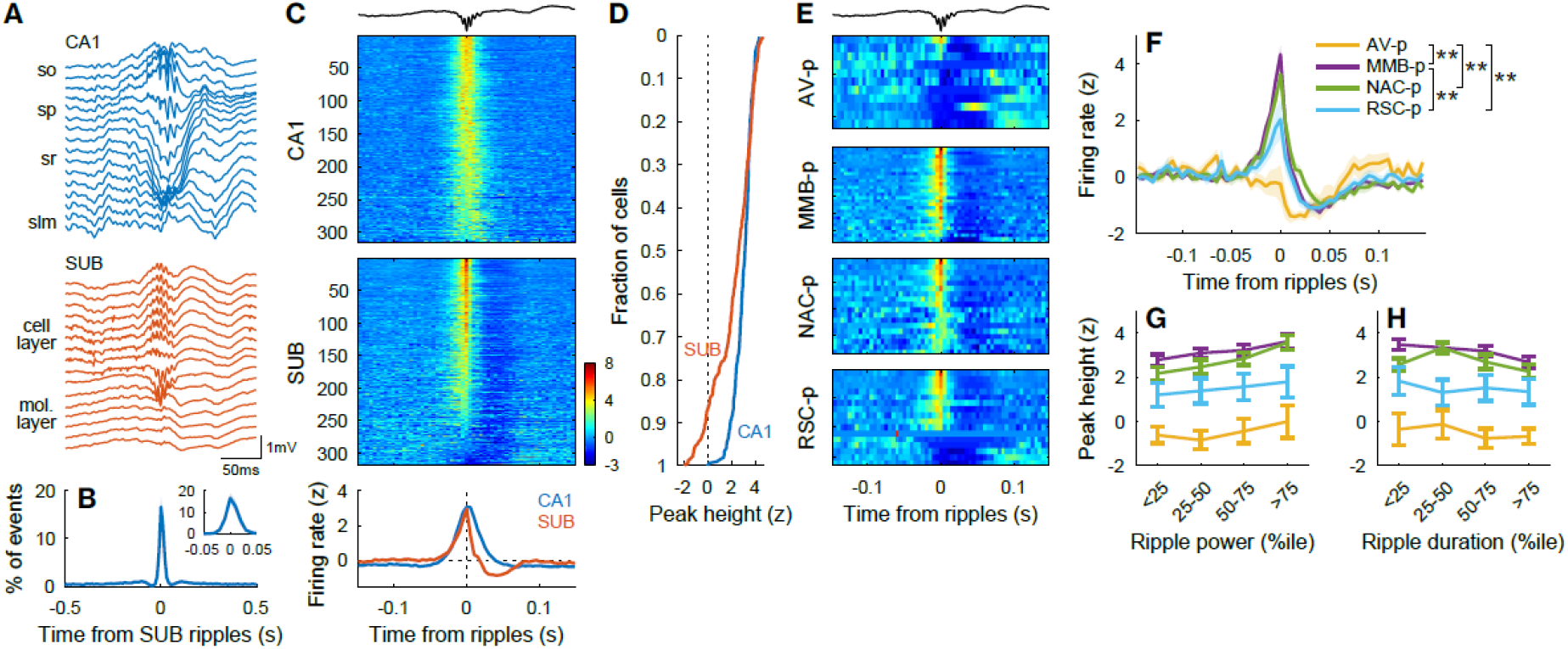
Projection-specific firing modulation by SPW-Rs during SWS. (A) Wide-band local field potentials (LFPs) simultaneously recorded in the CA1 (top) and SUB (bottom) showing SPW-Rs. (B) Cross-correlogram (CCG) of ripple events detected in the CA1 and SUB cell layers. Inset: the same CCG at a finer temporal scale. Mean (solid line) ± SD (shaded area). (C) Ripple-triggered average of firing rates (z-scored) of CA1 (top) and SUB (middle) neurons, and the firing rates averaged over CA1 or SUB neurons (bottom). Ripple events detected in single recording sites at the center of the SUB cell layer were used as the reference in (C–H). (C, E) Topmost trace: example SUB LFP showing SPW-Rs. (D) Mean z-scored firing rate averaged around the negative ripple peaks (–10 to 10 ms). (E) Ripple-triggered average of z-scored firing rates of individual SUB projection neurons. (F) Ripple-triggered average of z-scored firing rates of SUB projection neurons. ** *P* < 0.01, Tukey test for peak height. Mean (solid lines) ± SEM (shaded areas). (G, H) Peak height of firing rate (z-scored) as a function of ripple power (G) and ripple duration (H). The plot colors are the same as in (F). Mean ± SEM.

## Discussion

Using large-scale recordings combined with optogenetic identification of neuronal projections, three main findings were noted. Firstly, SUB neurons robustly represented multiple types of navigation-associated information as accurately as or even more accurately than CA1 neurons. Secondly, according to the type of information, SUB projection neurons selectively or uniformly distributed information to their target areas. Finally, information transmission of SUB projection neurons was distinctly controlled by theta oscillations and SPW-Rs in a target region-specific manner.

A key feature of subicular representations was that SUB neurons conveyed comparable (*i.e*., place and trajectory) or even greater (*i.e*., speed) information per unit time than that conveyed by CA1 neurons despite their broader tuning. This suggests that the sharply tuned, low firing-rate representation in the CA1 area is converted into a broadly tuned, high firing-rate representation in the SUB without losing the information content. The richer speed information in the SUB might be due to the integration of CA1 and entorhinal speed signals (*27, 28*). Moreover, the high firing rates in the SUB support the noise-resistant coding, which may be suitable for long-range projections suffering from substantial noises (*26*). These coding strategies were common across all types of information examined in this study. Thus, we propose that a key role of the SUB is to generate noise-resistant, accurate representations for multiple types of navigation-associated information. The coding strategy of constructing the high firing rate, noise-resistant representation may apply not only for the SUB but also for the entire hippocampal circuit. Indeed, the mean firing rates of principal cells progressively increase as they pass through the feed-forward network of the hippocampal formation (*i.e*., dentate gyrus → CA3 → CA1 → SUB) (*13, 32, 33*). Thus, one of the roles of the hippocampal circuit might be to convert the sparse firing in the dentate gyrus (*33, 34*) to progressively dense and robust representations.

We found that SUB projection neurons distribute information uniformly (*i.e*., place) or selectively (*i.e*., speed and trajectory) to downstream targets. Speed information prominently routed to the RSC may interact with visual-locomotion integration in this area (*35*). Trajectory information routed to the NAC may help goal-directed action to obtain a reward, which is thought to be the major function of NAC (*36*). Place information may be utilized in all target areas; particularly, AV, MMB, and RSC are all implicated in spatial learning (*37*). SUB projection neurons showed different preferred spike timing during theta oscillations, indicating that distinct information is sent out sequentially to different target areas in each theta cycle. SUB projection neurons were also distinctly activated or suppressed during ripples. Due to the widespread projections of the SUB, ripples broadcasted from the SUB might organize the brain-wide activity (*38*) through the modulation of downstream areas (*31, 39*). Exactly how the distributed information from the SUB is utilized in the target areas remains to be elucidated. Further studies are warranted to investigate the impact of SUB activity on information processing in the downstream areas.

## Acknowledgments

We thank Dr. Takaichi Fukuda for kindly providing the antibody to NOS, Dr. Hiroyuki Miyawaki for valuable suggestions on the manuscript, and Osaka City University Research Support Platform for allowing to use the confocal microscopy.

## Funding

This work was supported by JSPS KAKENHI (20H03356, 19H05225, 18H05137, 17K19462, 16H04656 and 16H01279) (K.M.), (20K06878, 19H04937, 17H05977, 17H05575 and 17K14939) (T.K.), JST PRESTO (JPMJPR1882) (T.K.), Toray Science Foundation (K.M.), Takeda Science Foundation (K.M. and T.K.), The Uehara Memorial Foundation (K.M. and T.K.), The Naito Foundation (K.M. and T.K.), The Nakajima Foundation (T.K.), SEI Group CSR Foundation (T.K.), Osaka City University Strategic Research Grant for young researches (T.K.), and Osaka City University Strategic Research Grant for basic researches (K.M.).

## Author contributions

T.K. and K.M. conceived the project. T.K. performed the experiments and analyzed the data except the retrograde tracing. R.U. performed the retrograde tracing and its analysis. T.K. and K.M. wrote the manuscript with inputs from all authors.

## Competing interests

The authors declare no competing interests.

## Data availability

Data is available upon reasonable request.

## Supplementary materials

Materials and Methods

Figures S1-S14

## Notes

### Competing Interest Statement

The authors have declared no competing interest.

