## Supplementary materials for "Robust information routing by dorsal subiculum neurons"

**This PDF file includes:**

Materials and Methods
Figs. S1 to S14

### Materials and Methods

All procedures of animal care and use were approved by the Institutional Animal Care and Use Committee of Osaka City University and were performed in accordance with the National Institutes of Health *Guide for the Care and Use of Laboratory Animals*.

### Anatomical tracing

AAVs for anatomical tracing were produced by co-transfecting plasmids into HEK293T cells, purified with heparin columns (HiTrap, GE Healthcare), and titered via qPCR, as described previously (*22*, *40*). Plasmids used for AAV production were as follows: pAAV-EF1α-EGFP (*22*), pAAV-EF1α-DIO-EYFP (No. 20296, Addgene), pAAV-pgk-Cre (No. 24593, Addgene), pXR1 (National Gene Vector Biorepository), pAAV2/6 (PL-T-PV0004, Penn Vector Core), pAAV-RC (AAV Helper-Free System, Stratagene), and pHelper (AAV Helper-Free System).

Stereotaxic injections were performed on male Long-Evans rats (n = 2, 13, 12 rats for anterograde tracing, retrograde tracing, and projection-specific labeling, respectively; 262–382 g; 7.9–10 weeks old on the day of surgery; SLC, Japan) under ~2% isoflurane anesthesia. For anterograde tracing, 400 nL of AAV1/2-EF1α-EGFP [$2.9 \times 10^{13}$ genome copies (GC)/mL] were injected into the left dorsal SUB [anterior-posterior from bregma (AP): –6.1 mm, mediolateral from the midline (ML): 3.0 mm, dorsoventral from the cortical surface (DV): 2.8 mm]. For retrograde tracing, either 400 or 1,200 nL/site of 0.5% w/v CTB488 (C34775, Thermo Fisher) were injected into one of the following brain areas: bilateral NAC (n = 4 rats; AP: +2.4 mm, ML: ±1.8 mm, DV: 6.4 mm), bilateral AV (n = 2 rats; AP: –2.0 or –1.7 mm, ML: ±1.6 mm, DV: 5.2 mm), ITN (n = 2 rats; AP: –2.0 mm, ML: 1.0 mm, DV: 6.6 mm, inserted from the right hemisphere at an angle of 10º in the coronal plane with the tip pointing towards the medial direction), bilateral RSC (n = 2 rats; AP: –3.8 mm, ML: ±0.4 mm, DV: 1.6 mm), and MMB (n = 3 rats; AP: –4.7, –4.2, or –3.2 mm, ML: 1.5 mm, DV: 8.9, 9.1, or 9.3 mm, at a 10º angle as described above). For projection-specific labeling (*22*), 1,000 nL/site of AAV1/2-EF1α-DIO-EYFP ($9.2 \times 10^{11}$ GC/mL) were injected into the bilateral dorsal SUB (AP: –6.1 mm, ML: ±3.0 mm, DV: 2.8 mm) and 400 nL/site of AAV6-pgk-Cre ($2.4 \times 10^{12}$ GC/mL) were injected into one of the following brain areas: bilateral NAC (n = 3 rats; AP: +2.4 mm, ML: ±1.8 mm, DV: 6.4 mm), bilateral AV (n = 2 rats; AP: –2.0 mm, ML: ±1.6 mm, DV: 5.2 mm), ITN (n = 2 rats; AP: –2.0 mm, ML: 1.0 mm, DV: 6.4 mm, at a 10º angle as described above), bilateral RSC (n = 3 rats; AP: –3.8 mm, ML: ±0.4 mm, DV: 2.0 mm), and MMB (n = 2 rats; AP: –4.7 mm, ML: 1.5 mm, DV: 8.9 mm, at a 10º angle as described above).

Immunohistochemistry was performed as described in **Histology**. Cells positive for CTB488, NOS, or PCP4 were counted manually using Fiji software at four or five AP levels of the SUB (350–2,150 µm posterior to the posterior commissure). In individual images, 10 × 2 and 10 × 5 grids were placed on the deep cell layer, rich in PCP4-positive cells (*41*), and the superficial cell layer, scarce in PCP4-positive cells, of the SUB, respectively. The density of CTB488-positive cells in each grid was obtained as the number of CTB488-positive cells in the grid divided by the grid area. Dendritic spines of EYFP-labeled dendrites were semi-automatically detected and measured from confocal image stacks (voxel size, $0.1 \times 0.1 \times 0.5\ \mu m^3$) using the NeuronStudio software as described previously (*42*). Spines were classified into three groups according to their shape: mushroom, thin, and stubby. Spine density was obtained as the number of spines divided by the dendrite length.

**Materials for extracellular recordings**

We used three types of 256-channel silicon probes (NeuroNexus). All types had eight shanks, with each shank containing linearly-aligned 32 recording sites (160–177 $\mu m^2$/site, 0.61 ± 0.23 MΩ at 1 kHz) but differing in their horizontal shank separation and vertical site spacing. Buzsaki256 probes had 300-µm shank separation and 50-µm vertical site spacing. A8x32-5mm-35-300-160 probes had 300-µm shank separation and 35-µm site spacing. A8x32-Edge-5mm-25-200-177 probes had 200-µm shank separation and 25-µm site spacing. The silicon probes were mounted on a 3D-printed microdrive with a movable screw (R0090B500, J.I. Morris), with which the silicon probe was gradually lowered to the SUB/CA1 area after implantation. For 8 of the 11 implantations, all recording sites were coated with poly(3,4-ethylenedioxythiophene) conducting polymer with a 1-µA direct current for 3 s (nanoZ, White Matter) to lower the impedance (0.44 ± 0.20 MΩ at 1 kHz after coating).

Laser diodes (450-nm light; PL450B, Osram) were coupled with a short optic fiber of 105-µm (FG105LCA, Thorlabs) or 200-µm (FP200URT, Thorlabs) core diameter (3.0–22.0 mm length depending on the target brain area). Light emission was controlled by a laser diode driver (LD202C, Thorlabs) and was measured before surgery from the tip of the optic fibers to reach 24–31 mW (with FG105LCA) and 42–88 mW (with FP200URT) with 100% laser output power.

**Surgery for recordings**

Eleven male Long-Evans rats (293–490 g; 8.9–15.3 weeks old on the day of surgery; SLC) were used. Under 1.6–2.0% isoflurane anesthesia, eight rats were stereotaxically implanted with four laser diode-coupled optic fibers targeting the left NAC (AP: +2.4 mm, ML: 1.8 mm, DV: 6.3 mm), left AV (AP: –2.0 mm, ML: 1.6 mm, DV: 4.6–4.8 mm), left RSC (AP: –3.8 mm, ML: 0.4 mm, DV: 1.5 mm), and MMB (AP: –4.7 mm, ML: 1.5 mm, DV: 8.7 mm, inserted from the right hemisphere at an angle of 10 degrees in the coronal plane with the tip pointing towards the medial direction), and were then injected with 800 nL/site of AAV1-hSyn-hChR2(H134R)-EYFP ($2.3 \times 10^{13}$ GC/mL; Penn Vector Core; diluted 1:1–1:5 before injection) using a pulled glass pipet (G-1, Narishige) into two sites in the left dorsal SUB (AP: –5.7 mm, ML: 2.2–2.5 mm, DV: 2.9–3.0 mm; and AP: –6.1 mm, ML: 3.2–3.5 mm, DV: 2.9–3.0 mm). Rats were finally implanted with a silicon probe targeting above the left dorsal SUB and distal CA1 area (center of the eight shanks at AP: –5.9 to –6.1 mm, ML: 2.7–3.3 mm, DV: 2.4 mm, with shanks parallel to the coronal plane). Two stainless steel screws (B000FN0J58, Antrin Miniature Specialties) inserted above the cerebellum served as indifferent and ground electrodes. A faraday cage connected to the ground electrode was placed around the implants. One rat was subjected to the same surgery, but the optic fiber implantation to the MMB was omitted for a technical reason. Two rats were implanted only with a silicon probe above the dorsal SUB and distal CA1 area (center of shanks at AP: –5.9 mm, ML: 2.8–3.3 mm, DV: 2.4 mm). All rats were housed individually post surgery.

**Data collection**

Electrophysiological data from behaving rats were acquired using a 256-channel, multiplexed recording system (KJE-1001, Amplipex) during behavioral tasks and sleep. Neurophysiological signals were amplified on a pre-amplifier module (HS-10, Amplipex) and acquired continuously at 20 kHz with 16-bit resolution. The silicon probe was lowered daily toward the SUB/CA1 area until large-amplitude units appeared approximately at the center of vertically-aligned recording sites at a depth of 2.7–3.8 mm from the cortical surface. Characteristic features (power, phase, and polarity) of theta oscillations and SPW-Rs were used as additional guides to determine the approximate locations of the recording sites relative to the SUB/CA1 cell layer (*6, 8*). The animal's position was tracked by monitoring two small light-emitting diodes (green and red, 5-cm separation) mounted above the head-stage using an overhead camera (c930e, Logicool) at ~30-Hz sampling rate. Single camera pixels corresponded to 0.41 cm for linear track and 0.52 cm for an open field, T-maze, zigzag maze, and rest sessions. LED positions were extracted and resampled to 39.0625 Hz for further analysis.

After the recording, to identify the location of recording sites, small electric lesions were made by passing anodal DC current (3 µA for 10 sec; A365, World Precision Instruments) through the most dorsal and ventral recording sites of each shank under anesthesia. Immunohistochemistry was performed as described in **Histology**. The tracks of silicon probe shanks and optic fibers were reconstructed from the images of serial sections.

The acquired LFP signals were down-sampled to 1,250 Hz for further analysis. Positive polarity is plotted upwards throughout the study. Spike sorting was first performed automatically using the Kilosort software (https://github.com/cortex-lab/KiloSort, https://github.com/MouseLand/Kilosort2), and then, clusters were adjusted manually using the phy gui (https://github.com/cortex-lab/phy), according to the developers' instructions. We measured cluster quality via the isolation distance and inter-spike interval (ISI) index, defined as the number of ISIs less than 2 ms divided by the number of ISIs between 2 ms and 10 ms multiplied by 4 (*40, 43*). Units satisfying all the following criteria were included for further analysis: isolation distance >20, ISI index <0.2, trough-to-peak amplitude >50 µV, and overall mean firing rate >0.1 Hz. Although we set these stringent criteria for unit quality, we cannot exclude that some units recorded in different sessions could have been identical, since spikes from sessions recorded on different days were clustered separately.

### Behavioral procedures

Rats were trained to perform four types of water-rewarded spatial tasks, namely, open field, linear track, alternating T-maze, and zig-zag maze, daily for 7–9 days before and 4–8 days after surgery. During training, the experimenter gave water drops randomly in the arena (for open field task) or at the reward ports (for the other tasks, see below) to motivate animal movement. In the alternating T-maze task, only correct trials were rewarded. Then, 13–22 days after surgery (6–13 days for two rats not injected with AAV), the final recording sessions were carried out for 2–3 consecutive days, with the four behavioral tasks performed each day for 20 min each and inter-task rest sessions performed for 40–80 min each. Rest sessions were also carried out before the first behavioral task and after the final task. On the final recording days, optical stimulation sessions to identify axonal projections were performed once (after all recording sessions of the day; one of the nine rats) or thrice (before, in the middle of, and after the recording sessions of the day; 8 of the 9 rats) while rats were resting in a small enclosure. During the periods of the behavioral experiments, rats were water-deprived in a way to maintain ~90% of their free-feeding body weight. All behavioral experiments were performed during the light period of the 12-h light/dark cycle.

The open-field task was performed using a square black arena (118 × 118 cm, 40-cm deep, with an A4-sized white cue card on one of the walls), in which rats freely foraged for randomly dispersed drops of water.

The linear track task was performed on an elevated black linear track (234-cm long, 6.5-cm wide, 54-cm high) on which rats were required to run back and forth to receive a 30-µL water drop supplied alternately at the edges of the track.

The alternating T-maze task was performed in a square black arena (118 × 118 cm, 40-cm deep) consisting of a start box (30 × 10 cm), central runway (stem, 98-cm long, 10-cm wide), and left/right arms (10-cm wide). Rats were enclosed by doors in the start box for ~8 s before allowed to run through the stem and arm. Rats were subjected to trials continuously and had to choose the other arm from that chosen in the previous trial to be rewarded with a 30-µl water drop at the end of the arm.

The zig-zag maze task was performed in a square black arena (118 × 118 cm, 40-cm deep), in which seven additional partitions (101-cm long, 30-cm high) were placed to form eight runways (118-cm long, 14-cm wide) connected with hairpin curves. Rats were rewarded with an 80-µl water drop alternately placed at the edges of the entire runway.

During the rest sessions, rats were placed in a small black enclosure (18 × 18 cm, 40-cm deep) not containing a water reward; optical stimulation to identify axonal projections was also performed in this enclosure.

### Optogenetic identification of projection targets

To identify the projection targets of recorded cells with antidromic spikes (*20*), we sequentially irradiated blue light pulses to the projection targets of the SUB while recording from the SUB and CA1. A single stimulation train consisted of 200 light pulses (10 pulses at 5 Hz repeated for 20 times with ~1-s intervals) and was repeated for combinations of 1- or 5-ms pulse durations with 12.5, 25, 50, 75, or 100% laser output power. Following these stimulation trains, closed-loop stimulation was performed to efficiently detect spike collisions. For this, spikes were detected online by high-pass filtering the extracellular signals at 300 Hz and by thresholding at 3 × root mean square, using a multi I/O processor (RX8, Tucker-Davis Technologies). Triggered by the detected spikes, a single light pulse (1-ms duration, 100% laser output power) was delivered to the target region with 1.8-ms delay. To avoid burst stimulation, the minimum inter-stimulation interval was set to 200 ms.

We identified projection neurons offline as follows. The peri-stimulus time histogram (0.5-ms bins) of individual cells was constructed for each stimulation train. Spikes in the first peak bin after stimulation onset (≥10 spikes and >mean + 3 SD of the pre-stimulus baseline period) and in its contiguous bins (>mean + 1 SD of the baseline period) were regarded as optically evoked spikes. Stimulation trains that gave low waveform correlation between the evoked and spontaneous spikes ($r < 0.9$) were excluded from further analysis. Fidelity for each stimulation train was calculated as the number of evoked spikes divided by that of stimulation trials (*i.e.*, 200). Jitter was defined as the SD of time from stimulation onset to the evoked spikes. Latency was defined as the mean time from stimulation onset to the evoked spikes. In the stimulation train with the highest fidelity, the units satisfying all following criteria were defined as the neurons projecting to the stimulated area: fidelity > 20%, jitter < 0.5 ms, and latency < 25 ms. These criteria were determined based on the units that passed the spike collision test (see below).

Spike collision test was performed as follows. For a given evoked spike latency $t$ ms, optical stimulation trials were divided into two types: collision trials that had one or more spontaneous spikes between $-t + 1$ ms and $t - 1$ ms (*i.e.*, shortly before the expected evoked spike latency $t$) and the rest of trials (non-collision trials). If the fidelity of optical spike generation in the collision trials was significantly less than that in non-collision trials ($P < 0.05$, $\chi^2$ test), the unit was regarded to pass the collision test. We used units that passed the spike collision test to define the criteria to identify projection target areas, so that 80% of the test-passed units were included in the criteria described above (fidelity > 20%, jitter < 0.5 ms, latency < 25 ms) (Fig. S7).

### Histology

At specific days after the surgery for anatomical tracing (anterograde tracing: 15 days, retrograde tracing: 7 days, projection-specific labeling: 20–21 days), or soon after the electric lesions for localizing silicon probes, rats were transcardially perfused with 0.9% saline, followed by 4% paraformaldehyde in 0.1 M phosphate buffer. The implanted silicon probe was pulled out before the brain was removed from the skull. Brains were stored in the same fixative overnight at 4 ℃ and then sectioned using a vibratome (VT1200S, Leica) at a 50-µm thickness parallel to the coronal plane. Sections were incubated sequentially with 5% bovine serum albumin (BSA) / 0.3% Triton X-100 in phosphate-buffered saline (PBS) for 30 min at room temperature, primary antibodies in 5% BSA/PBS overnight at 4 ℃, and the corresponding secondary antibodies conjugated with Alexa Fluor dyes in 5% BSA/PBS for either two hours at room temperature or overnight at 4 ℃. Some sections were counterstained with DAPI (0.5 µg/mL; D1306, Thermo Fisher) and/or NeuroTrace Red fluorescent Nissl (1:200; N21482, Thermo Fisher). Between the incubation procedures, sections were washed with PBS. The primary antibodies used were as follows: chicken anti-GFP (1:2,000; ab13970, Abcam), mouse anti-Cre recombinase (1:2,000; MAB3120, Millipore), mouse anti-NeuN (1:2,000; MAB377, Millipore), rabbit anti-PCP4 (1:200; HPA005792, Sigma), and sheep anti-NOS (1:10,000; gift from Dr. Takaichi Fukuda, originally from Dr. Piers C Emson) (*41*). The following secondary antibodies were used at 1:800 dilution: goat anti-chicken IgY conjugated with Alexa Fluor 488 (A-11039, Thermo Fisher), goat anti-mouse IgG with Alexa Fluor 594 (A-11032, Thermo Fisher), donkey anti-mouse IgG with Alexa Fluor 405 (ab175659, Abcam), donkey anti-rabbit IgG with Alexa Fluor 594 (A-21207, Thermo Fisher), and donkey anti-sheep IgG with Alexa Fluor 647 (A-21448, Invitrogen). Stained sections were mounted on coverslips with antifade mountant (P36961, Thermo Fisher). Tiled fluorescent images were taken using a confocal microscope (LSM700, Zeiss) equipped with a 10× [numerical aperture (NA) = 0.45, for anterograde tracing, and for localizing silicon probe and optic fibers], 20× (NA = 0.8, for retrograde tracing), and 63× (NA = 1.2, for dendritic spines) objectives.

### Analysis

#### *Cell classification*

We recorded a total of 791 well-isolated, large-amplitude units. Of these, 353 and 339 units were histologically identified to be localized in the CA1 area and SUB, respectively. Putative CA1 principal cells were defined as units that had >0.4-ms trough-to-peak spike width and <10-Hz overall mean firing rate. Putative SUB principal cells were defined as units that had >0.4-ms trough-to-peak spike width. The threshold of the mean firing rate was omitted when classifying SUB principal cells because a proportion of these cells shows high firing rates (Fig. S6)(*14*). Accordingly, 315 and 319 units were identified as CA1 principal cells and SUB principal cells, respectively.

We verified the criteria of principal cell classification of CA1 and SUB neurons by detecting putative monosynaptic interactions (*44*). Cross-correlograms (CCGs) of spike timing were constructed for all pairs of simultaneously recorded cells (0.1-ms bins, Gaussian smoothing with 0.5-ms σ). For each pair, 199 surrogate CCGs were also constructed from data with random jittering of spike timing by –5 to 5 ms in one of the paired cells. The 99% global band significance level was calculated as described previously (*44*). CCGs with a peak or trough crossing the 99% significance level between 0 ms and 5 ms were taken as the candidate pairs of monosynaptic excitation or inhibition, respectively. Of these, dull peaks and troughs, unlikely to be due to monosynaptic spike transmission and suppression, were excluded by visual inspection. Nearly all CCG-based putative excitatory cells (CA1: 44 of 45 neurons, SUB: 40 of 40 neurons) were confirmed to meet the criteria of principal cells described above, and all CCG-based putative inhibitory cells (CA1: 6 of 6 neurons, SUB: 5 of 5 neurons) had <0.4-ms spike width (Fig. S6). Hereafter, we analyzed only the CA1 and SUB principal cells.

*Place representation on a linear track*

We examined the spatial firing patterns on a linear track by constructing one-dimensional rate maps for individual cells. A rate map consisted of firing rates at position bins (2-cm long, excluding the edges of the track), calculated by dividing the number of spikes in each position bin by the duration spent in that bin, after having individually smoothed both the numerator and denominator with a Gaussian filter (σ = 2 cm). For each cell, rate maps were constructed separately for eastbound and westbound trials, and the rate map with the higher peak firing rate (*i.e.*, the rate map of the cell's preferred direction) was used for further analysis. Cells with peak firing rates less than 1 Hz in the preferred direction were excluded from the analysis. The mean rate was defined as the mean firing rate during trials in the preferred direction. Spatial information per spike ($I_{spike}$, bits/spike) and spatial information per second ($I_{sec}$, bits/s) were calculated by

$$I_{spike} = \sum_i p_i \frac{\lambda_i}{\lambda} log_2\left(\frac{\lambda_i}{\lambda}\right)$$

$$I_{sec} = \sum_i p_i \lambda_i log_2\left(\frac{\lambda_i}{\lambda}\right)$$

where $p_i$ is the proportion of time spent in the *i*-th position bin, $\lambda_i$ is the firing rate in the *i*-th position bin, and $\lambda$ is the mean firing rate during trials (*23*).

The chance level of $I_{sec}$ was obtained by a trial-wise shuffling procedure, by which the rate map of every trial was circularly shifted relative to the track position by a random interval, with the end of the trial wrapped to the beginning before obtaining the rate map averaged over trials. The shuffling was repeated 1,000 times for each cell. A cell was defined as a place cell if its $I_{sec}$ exceeded the 99th percentile of the $I_{sec}$ from the shuffled data obtained from all CA1 and SUB cells.

Mutual information (bits) of firing rates and places was estimated according to a previous study (*24*) by

$$Mutual\ information = \sum_i \sum_j p_{ij} log_2\left(\frac{p_{ij}}{p_i p_j}\right)$$

where $p_i$ is the probability that the animal is in the *i*-th position bin (bin size, 2 cm), $p_j$ is the probability that the instantaneous firing rate of a cell is in the *j*-th firing rate bin (four bins, see below), and $p_{ij}$ is the joint probability between the *i*-th position bin and *j*-th firing rate bin. The instantaneous firing rate of a cell was obtained by sorting the cell's spike timing into 25.6-ms bins, followed by Gaussian smoothing (σ = 128 ms) of the binned firing rate. The instantaneous firing rates were sorted into four firing rate bins with threshold values at 25, 50, and 75 percentiles of the non-zero instantaneous firing rate values.

*Place decoding on a linear track*

We performed memoryless Bayesian decoding of the rat's position to estimate the positional information conveyed by CA1 and SUB cell populations (*25*). The probability of the rat's position (pos) across M total position bins in time window τ (250 ms) containing neural spikes (spikes) was

$$\Pr(pos|spikes) = U / \sum_{i=1}^{M} U$$

where

$$U \; = \left( \prod_{j\,=\,1}^{N} f_j(pos)^{n_j} \right) exp\left( -\tau \sum_{j\,=\,1}^{N} f_j(pos) \right).$$

The $f_j$(pos) is the rate map of the $j$-th unit, $n_j$ is the number of spikes of the $j$-th unit in the time window, and $N$ is the number of units used for decoding. Decoding performance was estimated using leave-one-out cross-validation as follows. From a given set of trials in the recording session, a trial was chosen as test data and the rest of trials were used as training data. The decoding result, Pr(pos|spikes), was obtained using rate maps $f_j$(pos) constructed from the training data and spike train $n_j$ in the test data. The position with the highest Pr(pos|spikes) across the position bins was defined as the decoded position of the given time window. The decoding error of the test data was calculated as the mean Euclidean distance between the decoded and the observed positions. This procedure was repeated to assign every trial as test data, and the decoding errors from all test data were averaged to obtain the decoding error of the session. The chance-level distribution of the decoding error was estimated by a shuffling procedure repeated 100 times: The decoded positions were randomly permuted along time windows, and the decoding error was calculated.

Dependency of the decoding performance on the number of used cells was measured by random cell sub-sampling. Of the total simultaneously recorded CA1 or SUB cells ($N_{total}$), decoding was performed using a certain number of random cell subsets, and the decoding error was calculated. This procedure was repeated 100 times (or for all possible combinations of the cell group if the number of combinations was less than 100), and the obtained decoding errors were averaged over the repetition. By changing the number of units supplied for the decoding from 1 to $N_{total}$, we obtained the decoding error for each number of units.

To estimate the contribution of single units to the decoding performance, we performed a Jackknife procedure after random sub-sampling of the units. Of the total simultaneously recorded CA1 or SUB units, one unit was chosen as a test unit and nine units were randomly chosen from the remaining units. Recording sessions with less than 10 simultaneously recorded CA1 or SUB units were excluded from this analysis. Decoding was performed using the nine units with or without the test unit, and the decoding improvement of the test unit was calculated as the difference in the decoding error between the two decoding error values. The random choice of nine units was repeated 100 times from all possible combinations of nine units without replacement (or for all possible combinations of choices if the number of combinations was less than 100), and the decoding improvement of the test unit was averaged over the repetition. This procedure was further repeated to assign every recorded unit as a test unit once.

The robustness of the decoding performance against noises was estimated by randomly removing (up to 99% of the number of spikes of each unit) or adding spikes to every unit at a fixed frequency over the entire recording session before performing the decoding procedure with all simultaneously recorded CA1 or SUB cells. The removal/addition procedure was repeated 100 times to obtain the decoding error, averaged over the repetition. Recording sessions with less than five simultaneously recorded units were excluded from this analysis.

The entire decoding procedure was performed separately for eastbound and westbound runs, which were pooled later.

*Place representation in an open field*

We analyzed the spatial firing patterns in the open field similarly to those on the linear track. A rate map was constructed by 2 × 2 cm position bins using a two-dimensional Gaussian filter ($\sigma$ = 4 cm) for smoothing. Units with a mean firing rate less than 0.1 Hz in the open field were excluded from the analysis. Spatial information per spike ($I_{spike}$, bits/spike) and that per second ($I_{sec}$, bits/s) were calculated exactly as those on the linear track. A session-wise shuffling procedure was performed, by which the spike train of the whole open-field session was circularly time-shifted relative to the rat's position by a random interval between 30 s and the length of the session minus 30 s, with the end of the trial wrapped to the beginning.

We performed memoryless Bayesian decoding ($\tau$ = 1 s) as in the case of the linear track. Decoding performance was estimated using five-fold cross-validation as follows. The four-fifth periods of the recording session (*e.g.,* 0–960 s of the 1,200-s session) were chosen as the training data and the remaining periods (*e.g.,* 960–1,200 s) were assigned as the test data. The probability of the rat's position across position bins, Pr(pos|spikes), was calculated from rate maps $f_j$(pos) constructed from the training data and spike train $n_j$ in the test data, as described above. This procedure was repeated five times to decode from all periods of the recording session, and the decoding errors from the five periods were averaged to obtain the decoding error of the session. The chance-level distribution of the decoding error was estimated by a shuffling procedure (100 repetitions), by which the decoded positions were randomly permuted along time bins, and the decoding error was calculated from the shuffled decoded positions and the observed positions. Dependency of the decoding performance on the number of used cells and the robustness of the decoding performance against noises were also estimated as described for the linear track.

*Speed representation in an open field*

Speed representation in the open field was analyzed as described previously (*27*, *28*). Neuronal spikes were sorted into 25.6-ms time bins. The instantaneous firing rate was obtained by dividing the numbers of spikes for each cell by the bin size and smoothed with a Gaussian filter ($\sigma$ = 256 ms). Units with mean firing rate less than 0.1 Hz in the open field were excluded from the analysis. Each rat's positions were smoothed with a Gaussian filter ($\sigma$ = 256 ms). Instantaneous running speed was calculated by dividing the distance of the smoothed position between adjacent bins by the bin size (25.6 ms). Changing the SD of Gaussian filters to 128 or 512 ms for both firing rate and position gave similar results. Periods slower than 2 cm/s and faster than 50 cm/s were removed from the analysis. Speed scores were defined for each cell as the Pearson product-moment correlation coefficients between the cell's instantaneous firing rate and the rat's instantaneous running speed. We calculated speed information as follows (*27*). First, for each cell, the speed tuning curve of the firing rate against speed was constructed using bins of 4 cm/s, from 2 cm/s to 50 cm/s. Then, speed information per spike ($I_{spike}$, bits/spike) and speed information per second ($I_{sec}$, bits/s) were obtained by

$$I_{spike} = \sum_i p_i \frac{\lambda_i}{\lambda} log_2 \left(\frac{\lambda_i}{\lambda}\right)$$

$$I_{sec} = \sum_i p_i \lambda_i log_2 \left(\frac{\lambda_i}{\lambda}\right)$$

where $p_i$ is the proportion of time spent in the $i$-th speed bin, $\lambda_i$ is the firing rate in the $i$-th speed bin, and $\lambda$ is the mean firing rate in the open field.

Chance-level statistics were calculated by a shuffling procedure (100 repetition), by which the spike train was circularly time-shifted relative to the rat's position by a random interval between 30 s and the length of the session minus 30 s, with the end of the trial wrapped to the beginning. A cell was defined as a positive speed (p-speed) cell or negative speed (n-speed) cell if its speed score exceeded the 99th percentile or was lower than the 1st percentile, respectively, of the distribution of speed scores from the shuffled data from all CA1 and SUB principal cells.

*Decoding of running speed*

Decoding of running speed was performed using five-fold cross-validation according to a previous study (*28*). Of the 1,200-s recording session in the open field, one fifth of the period (*e.g.*, 0–240 s) was assigned as test data and the remaining four fifths (*e.g.*, 240–1,200 s) were used as training data. A linear relationship between the firing rate and running speed averaged over 1-s bins was expressed as

$$S_{tr} = R_{tr} f$$

where $S_{tr}$ is a column vector with the speed bins of the training data, $R_{tr}$ is a matrix containing the corresponding firing rate bins for each neuron as columns and an additional column of 1s to account for y intercepts, and $f$ is a linear filter used as a column vector with a length equal to the number of cells plus 1. The linear filter $f$ was obtained by

$$f = {R_{tr}}^{+} S_{tr}$$

where ${R_{tr}}^{+}$ is the Moore-Penrose pseudoinverse of $R_{tr}$. Once $f$ was obtained, the decoded speed $S_{dec}$ was calculated by

$$S_{dec} = R_{test} f$$

where $R_{test}$ is the firing rate matrix of the test data. Decoding accuracy was defined as the Pearson correlation coefficient between $S_{dec}$ and the observed speed $S_{test}$. The chance level of the decoding accuracy was estimated by a shuffling procedure repeated 100 times, by which $S_{dec}$ was time-shifted by a random interval between 30 s and 210 s relative to $S_{test}$. The decoding procedure was repeated five times to decode all periods of the recording session, and the obtained decoding accuracy was averaged over the five periods. Dependency of the decoding performance on the number of used cells was measured by random sub-sampling of the cells, as described above. The single-unit decoding accuracy refers to the accuracy obtained from the decoding using single units.

Robustness of the decoding performance against noises was estimated by randomly removing (up to 99% of the number of spikes of the unit) or adding spikes to every unit at a fixed frequency over the entire recording session before performing the decoding procedure with all the simultaneously recorded CA1 or SUB units. The removal/addition procedure was repeated 100 times to obtain the decoding accuracy, averaged over the repetition. The normalized accuracy was obtained as the decoding accuracy values divided by the corresponding decoding accuracy without the spike removal/addition. Recording sessions containing less than five units were excluded from this analysis.

*Trajectory-dependent firing in an alternating T-maze*

Trajectory-dependent firing in the start box of the T-maze was analyzed. A trial consisted of a wait period of ~8 s in the start box, running through the stem and left or right arm, reward acquisition at the end of the arm, and return to the start box. To estimate the strength of trajectory dependent firing in the start box, we obtained the following parameters for each neuron using firing rates in the start box: trajectory information per spike ($I_{spike}$, bits/spike) and trajectory information per second ($I_{sec}$, bits/s), calculated by

$$I_{spike} = \sum_i p_i \frac{\lambda_i}{\lambda} log_2\left(\frac{\lambda_i}{\lambda}\right)$$

$$I_{sec} = \sum_i p_i \lambda_i log_2\left(\frac{\lambda_i}{\lambda}\right)$$

where $i$ is the identifier of left or right choices, $p_i$ is the proportion of choice $i$, $\lambda_i$ is the mean firing rate in the start box for the subsequent choice $i$, and $\lambda$ is the mean firing rate in the start box. Units with $\lambda_i < 0.1$ Hz for both left and right choices were excluded from the analysis. The rate change ratio was defined as $|\lambda_L - \lambda_R|/\max(\lambda_L, \lambda_R)$, where $\lambda_L$ and $\lambda_R$ are the firing rates in the start box averaged over left- and right-arm trials, respectively. The auROC (ranging from 0 to 1) was calculated to estimate the goodness of binary classifier (*i.e.*, left- or right-arm trials) from the mean firing rates in the start box using the 'perfcurve' function in Matlab. For auROC values less than 0.5, 1 – auROC was assigned as the new auROC value, as the experimenter had no prior knowledge on the type of trials (left or right) that neurons showed a higher firing rate. We defined trajectory-dependent cells as neurons whose firing rates in the start box between left- and right-arm trials were significantly different ($P < 0.05$, Wilcoxon rank sum test).

*Decoding of trajectory*

Trajectory decoding was performed using a support vector machine algorithm and leave-one-out cross-validation according to a previous study (*29*). From a given set of trials in the recording session, one trial was assigned as test data and the remaining trials were used as the training dataset. A binary classifier was constructed to predict the next trajectory (left or right) from the firing rates in the start box, using 'fitcsvm' function in Matlab, in which the input arguments were matrix $F$ consisting of the firing rates of individual neurons (columns) in the training trials (rows) and vector $y$ consisting of the trajectory label (–1, 1) of the training trials. Using the firing rates in the test trial as an input to the constructed classifier, the trajectory in the test trial was predicted. This decoding procedure was repeated to assign every trial as test data once, and the decoding accuracy was defined as the proportion of correctly predicted trials. Dependency of the decoding performance on the number of used cells was measured by random sub-sampling of the cells, as described above. The chance level of the decoding accuracy was estimated by a shuffling procedure repeated 100 times, by which trajectory labels were randomly shuffled before constructing the classifier.

The robustness of the decoding performance against noises was estimated by randomly removing (up to 99% of the number of spikes of the unit) or adding spikes to every unit at a fixed frequency over the entire recording session before performing the decoding procedure with all the simultaneously recorded CA1 or SUB units. The removal/addition procedure was repeated 100 times to obtain the decoding accuracy averaged over the repetition. The normalized accuracy was obtained as follows: (decoding accuracy – 0.5) / (corresponding decoding accuracy without the spike removal/addition – 0.5). Recording sessions containing less than five units were excluded from this analysis.

*Behavioral states*

Throughout the recording sessions, each second of behavior was classified into four states: RUN, REST, REM sleep, or SWS. This classification was carried out by the visual inspection of power spectra of SUB LFPs, SUB raw LFP traces, and electromyogram (EMG)-related signals extracted as correlations of LFPs in distant channels after filtering at 300–600 Hz (*45*). RUN states were defined as the periods during behavioral tasks with prominent theta oscillations and EMG-related signals. REST was defined as the awake, resting state detected during rest sessions without theta oscillations but with EMG-related signals. REM sleep was detected by the presence of robust theta oscillations and the absence of EMG-related signals during rest sessions. SWS was detected by the absence of both theta oscillations and EMG-related signals during rest sessions.

*Theta oscillations*

Power spectra density of LFP during RUN and REM periods was obtained using Welch periodogram method (50% overlapping Hamming windows with a length of 2 s). Theta power was obtained as the mean power spectra density (in dB) between 5 and 10 Hz. A recording site with maximal theta power approximately at the center of the SUB cell layer was used as the reference site to determine phase deviation and phase locking. Instantaneous theta phase was derived from the Hilbert transform of the bandpass-filtered (5–10 Hz) LFP trace (peaks = 0, 360 degrees and troughs = 180 degrees throughout this study). Theta phase deviation was calculated as the mean circular distance of instantaneous theta phases between the reference recording site and the recording site of interest. Spike theta phase, *i.e.*, the theta phase at which the spike occurred, was obtained for every spike. The preferred theta phase of each neuron was defined as the circular mean of the spike theta phases. PPC, a firing rate-insensitive measure of phase locking strength, was calculated by

$$PPC = \frac{2}{N(N-1)} \sum_{i=1}^{N-1} \sum_{j=i+1}^{N} \left\{\cos(\theta_i)\cos(\theta_j) + \sin(\theta_i)\sin(\theta_j)\right\}$$

where $N$ is the total spike number, and $\theta_i$ is the spike theta phase of the $i$-th spike (*30*).

*Sharp-wave/ripple-associated firing*

Spike timing along SPW-Rs was examined during SWS and REST periods. Ripple-band LFP signal was obtained as the bandpass-filtered LFP signal (140–230 Hz) at the center of the SUB cell layer. Normalized ripple power was calculated as the z-scored moving average (window size, 11 samples) of squared ripple-band LFP signals. Periods with a normalized ripple power exceeding three were collected as candidate ripple events. Temporally close candidate events (<30-ms inter-event intervals) were merged into single events. The candidate events with low peak normalized ripple power (<7), too short duration (<15 ms), or too long duration (>300 ms) were discarded, and the remaining events were defined as ripple events. Ripple timing was determined as the timing of the negative peak of the ripple-band LFP signal in each event. Ripple power was defined as the peak value of the normalized ripple power in each ripple event. Ripple duration was defined as the duration of the normalized ripple power exceeding three. A peri-event time histogram (5-ms bins) was constructed for each ripple event for each cell, averaged across all events, and z-scored. The peak height for each neuron was defined as the mean z-scored peri-event time histogram between –10 to +10 ms. All data analyses were performed by custom-written MATLAB codes.

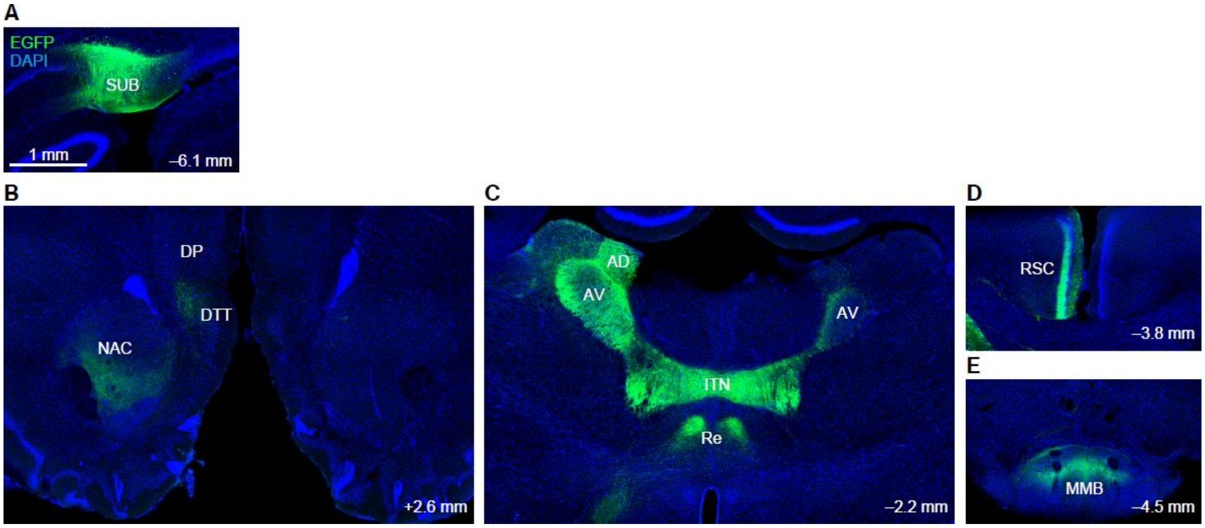


**Fig. S1. Anterograde tracing from the subiculum (SUB)**

We determined subicular projection targets by anterograde tracing. We injected adeno-associated virus (AAV) with chimeric serotype 1/2 expressing enhanced green fluorescent protein (EGFP) under the control of EF1α promoter (AAV1/2-EF1α-EGFP) into the left dorsal SUB (A) and investigated the distribution of EGFP-labeled axons throughout the brain (n = 2 rats). EGFP-labeled axons were observed in the ipsilateral nucleus accumbens (NAC), ipsilateral dorsal tenia tecta (DTT), ipsilateral dorsal peduncular cortex (DP) (B), bilateral anteroventral thalamic nucleus (AV), anterodorsal thalamic nucleus (AD), interanteromedial thalamic nucleus (ITN), nucleus reuniens (Re) (C), the superficial layer of the ipsilateral granular retrosplenial cortex (RSC) (D), and medial mammillary body (MMB) (E). Labeled axons were also observed in the entorhinal cortex (data not shown). All images are coronal sections. Numbers indicate an approximate distance of the sections from the Bregma.

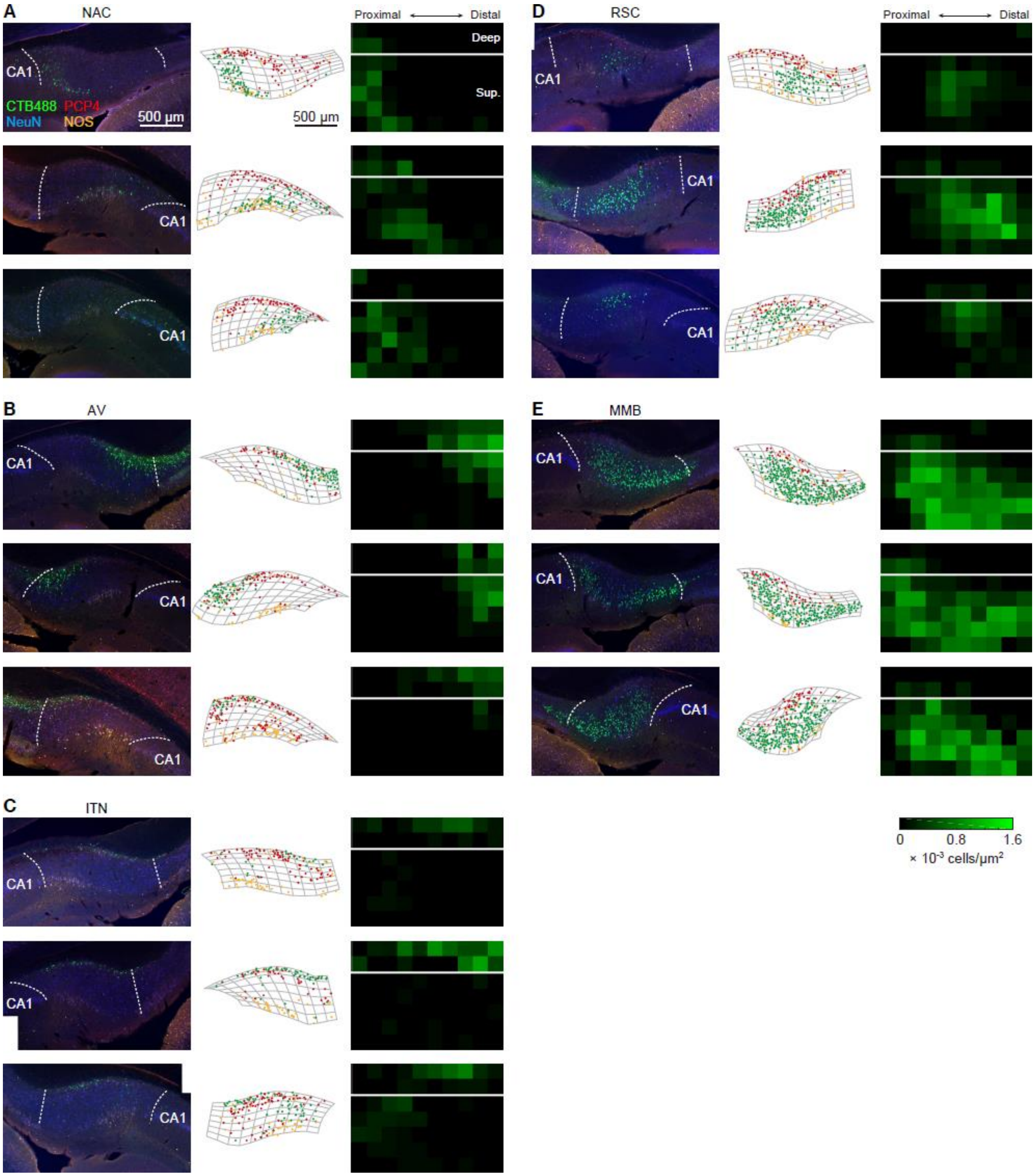


**Fig. S2. Additional examples of retrograde tracing**

(A–E) Example coronal images of the dorsal subiculum (SUB) containing retrogradely-labeled cholera toxin B conjugated with Alexa Fluor 488 (CTB488)-positive cells. We performed immunohistochemistry for Purkinje cell protein 4 (PCP4, rich in the SUB deep layer) and nitric oxidase synthase (NOS, rich in the proximal superficial part of the SUB cell layer), in order to visualize the laminar structure within the SUB cell layer (*41*). Three sections of different hemispheres are shown for each CTB488-injection site: nucleus accumbens (NAC; A), anteroventral thalamic nucleus (AV; B), interanteromedial thalamic nucleus (ITN) (C), retrosplenial cortex (RSC; D), and medial mammillary body (MMB; E). Left columns in (A–E) show confocal images of the SUB. Dotted lines represent borders of the SUB cell layer. Note that proximal-distal directions are flipped image-by-image (see the labels of the CA1 area), since images were taken from both left and right hemispheres. Middle columns represent the soma locations of detected CTB488-positive (green), PCP4-positive (red), and NOS-positive (orange) cells, overlaid on the segmented SUB cell layer (gray mesh). The deep layer (top two rows of each mesh) contains most PCP4-positive cells. The images in the middle columns have the same proximal-distal directions with the corresponding confocal images in the left column. Right columns show the density maps of CTB488-positive cells in deep (top, close to alveus) and superficial (bottom, close to molecular layer) SUB cell layers. All images in the right columns are aligned so that the proximal SUB is shown at the left side of the image. Note that the images shown at the top rows of (A–E) are also shown in Figure 1.

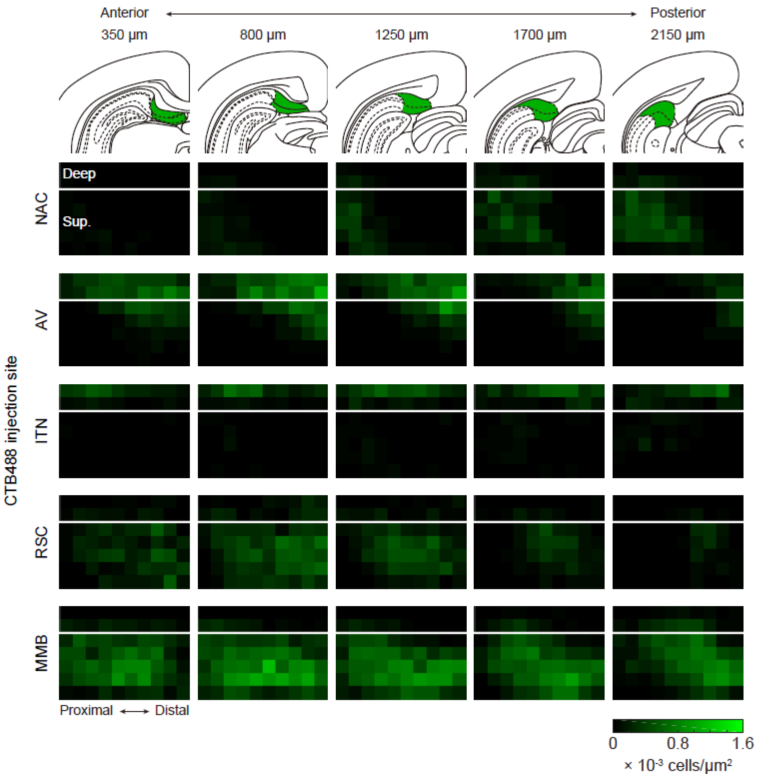


**Fig. S3 Population data of retrograde tracing**

Density maps of cholera toxin B conjugated with Alexa Fluor 488 (CTB488)-positive cells averaged across hemispheres for each injection site (rows) at each anterior-posterior distance from the posterior commissure (columns). Numbers indicate the distances of sections in the posterior direction from the posterior commissure. Corresponding brain atlas images (*46*) containing the subiculum (SUB; green) are shown below. Individual maps show CTB488-positive cell density in the SUB cell layer along proximal-distal (left-right) and deep-superficial (top-bottom) axes. Interanteromedial thalamic nucleus (ITN)-projecting neurons were primarily located at the deep SUB cell layer, while anteroventral thalamic nucleus (AV)-projecting neurons were located at the distal-deep cell layer. Nucleus accumbens (NAC)-projecting neurons were found at the proximal one-third of the SUB cell layer, while retrosplenial cortex (RSC)-projecting neurons showed complementary localization at the distal-superficial cell layer. Medial mammillary body (MMB)-projecting neurons were located at the superficial cell layer, containing few PCP4-positive cells. n = 6, 4, 4, 4, and 6 hemispheres for injections in the NAC, AV, ITN, RSC, and MMB, respectively.

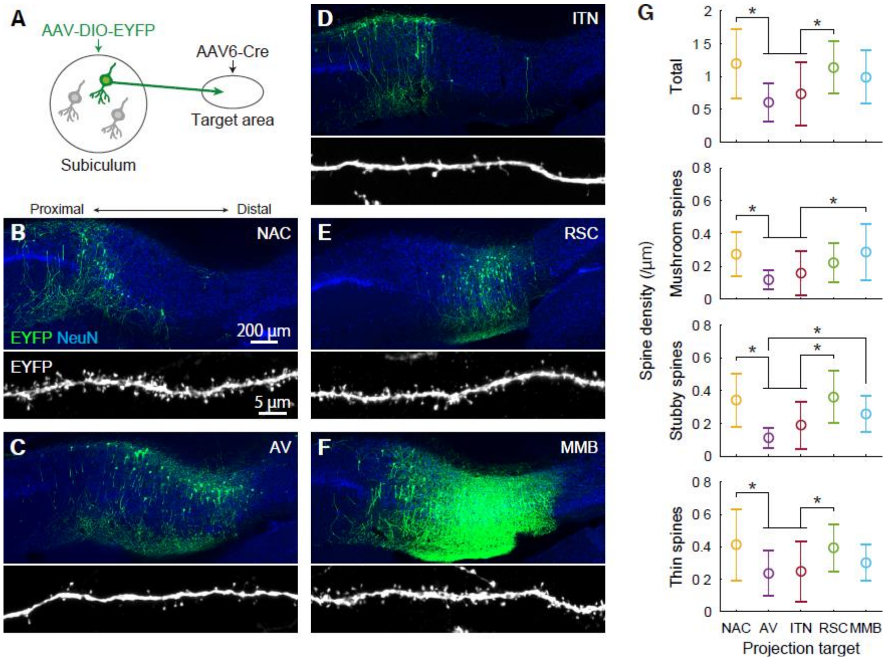


**Fig. S4. Dendritic morphology of projection neurons in the subiculum (SUB)**

(A) We visualized the fine morphology of SUB projection neurons targeting a specific area by injecting the retrograde-infecting AAV6-pgk-Cre (*i.e.*, serotype 6, expressing Cre recombinase under the control of pgk promoter) into the target area of interest and the Cre-dependent, EYFP-expressing AAV (AAV1/2-EF1α-DIO-EYFP) into the dorsal SUB.

(B–F) Representative images of coronal SUB sections containing EYFP-labeled projection neurons (top) and apical dendrites (bottom). Injection sites of AAV6-pgk-Cre included one of the following areas: nucleus accumbens (NAC; B), anteroventral thalamic nucleus (AV; C), interanteromedial thalamic nucleus (ITN; D), retrosplenial cortex (RSC; E), and medial mammillary body (MMB; F).

(G) Dendritic spine density per unit length of the apical dendrite. n = 17, 17, 25, 21, and 20 cells for NAC, AV, ITN, RSC, and MMB, respectively. * $P < 0.05$, Tukey test. Mean ± standard deviation.

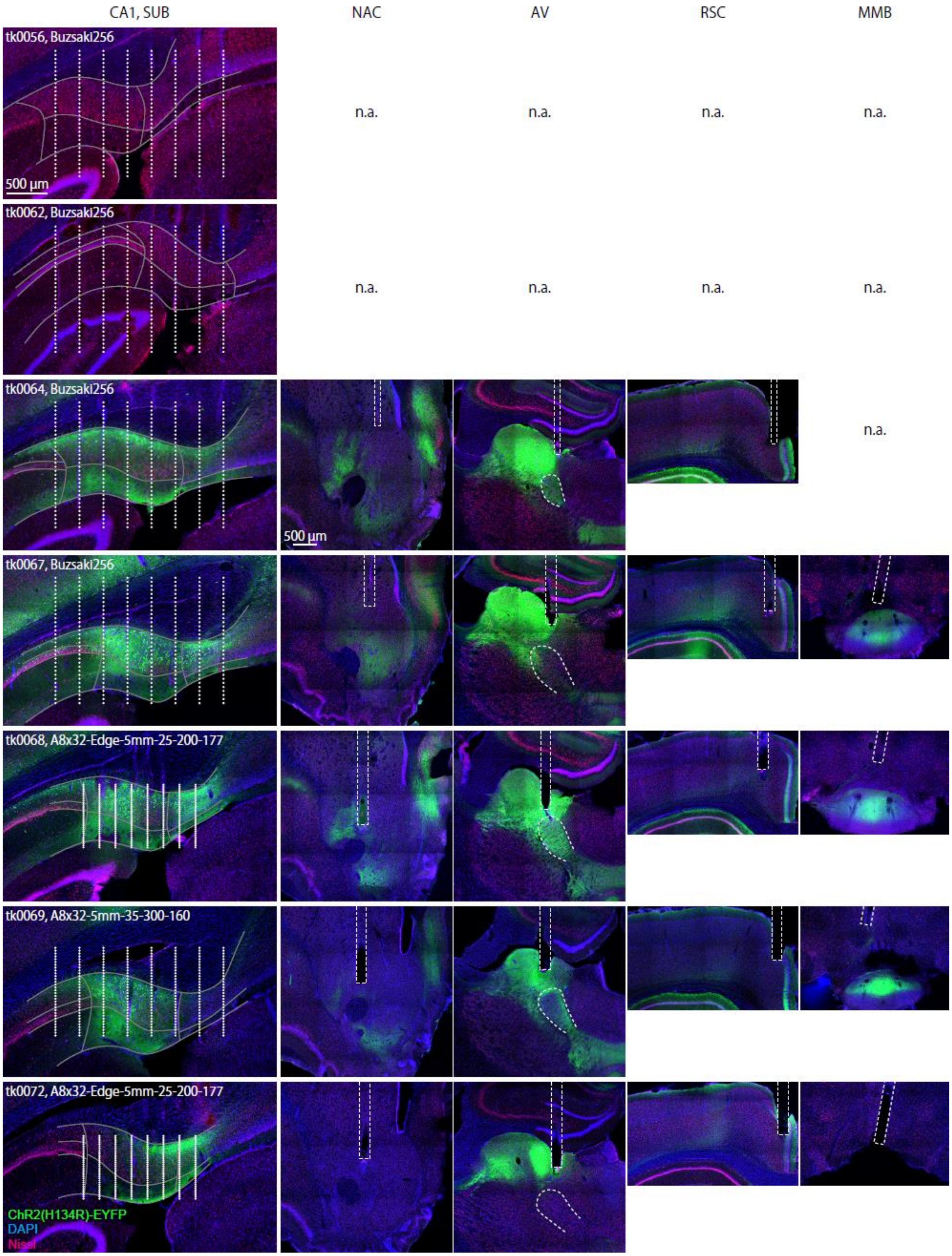

CA1, SUB
NAC
AV
RSC
MMB
tk0056, Buzsaki256
500 µm
n.a.
n.a.
n.a.
n.a.
tk0062, Buzsaki256
n.a.
n.a.
n.a.
n.a.
tk0064, Buzsaki256
500 µm
n.a.
tk0067, Buzsaki256
tk0068, A8x32-Edge-5mm-25-200-177
tk0069, A8x32-5mm-35-300-160
tk0072, A8x32-Edge-5mm-25-200-177
ChR2(H134R)-EYFP
DAPI
Nissl

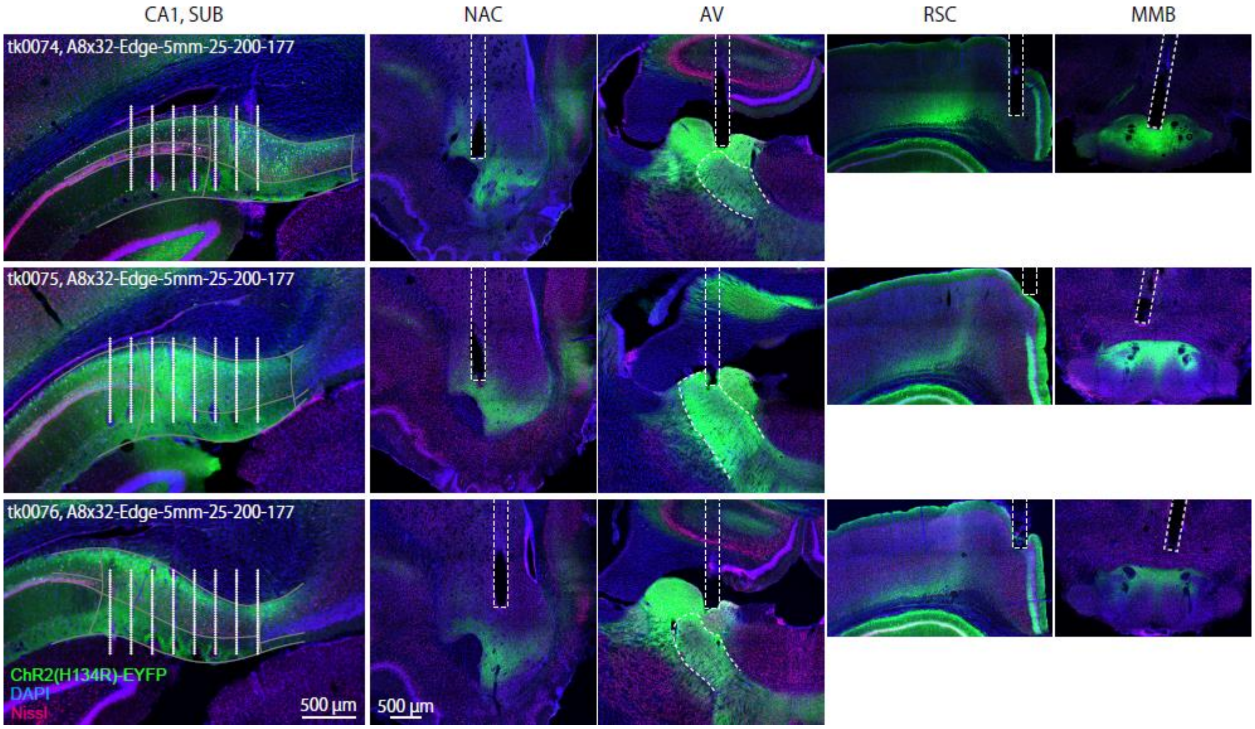


**Fig. S5. Post-hoc histology**

We confirmed the locations of recording sites, optical fibers, and channelrhodopsin-2 (ChR2) expression (green) by *post-hoc* histology. Sections were counterstained for Nissl (red) and DAPI (blue). All recorded rats except the one shown in Fig. 1 are shown in this figure. Text labels in the leftmost column show the animal ID and the implanted silicon probe. White dots represent the estimated recording sites. Gray lines are the anatomical borders of the CA1, subiculum (SUB), presubiculum, and their cell layers. Dotted rectangles in the 2nd to 5th columns show the locations of optical fibers. Dotted curves show the outline of the anteroventral thalamic nucleus (AV). For rats tk0056 and tk0062, no AAV injection or optic-fiber implantation was performed. For the rat tk0064, no optic fiber was implanted to the MMB. Note that, for clear visualization of the localization of EYFP-labeled SUB axons, the image brightness was linearly adjusted for individual images in this figure and thus, the image intensity should not be compared between images. NAC, nucleus accumbens; ITN, interanteromedial thalamic nucleus; RSC, retrosplenial cortex; MMB, medial mammillary body.

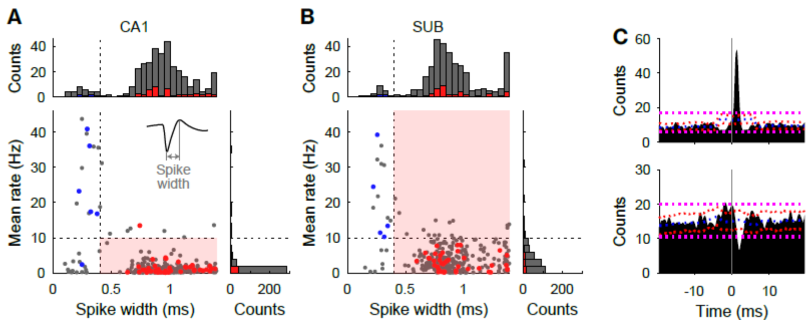


**Fig. S6. Cell type classification and verification**

(A, B) We classified putative principal cells based on spike width and overall mean firing rates. Putative CA1 principal cells (A) were defined as units with >0.4-ms spike width and <10-Hz mean firing rate (shaded area in the scatter plot). Putative subiculum (SUB) principal cells (B) were defined as units with >0.4-ms spike width (shaded area). The spike width was measured as the trough-to-peak width of the mean wide-band spike waveform (inset). Each dot represents single units. Among them, units that were verified to be putative excitatory and putative inhibitory by cross-correlogram analysis (C) are shown as red and blue, respectively.

(C) Representative cross-correlograms of spike timing between pairs of neurons. As described previously (*44*), a narrow peak (top) and a trough (bottom) with short latency identify a reference cell as being putatively excitatory and putatively inhibitory, respectively. Blue line: time-jittered mean; red line, point-wise comparison ($P < 0.01$); magenta line: global comparison ($P < 0.01$).

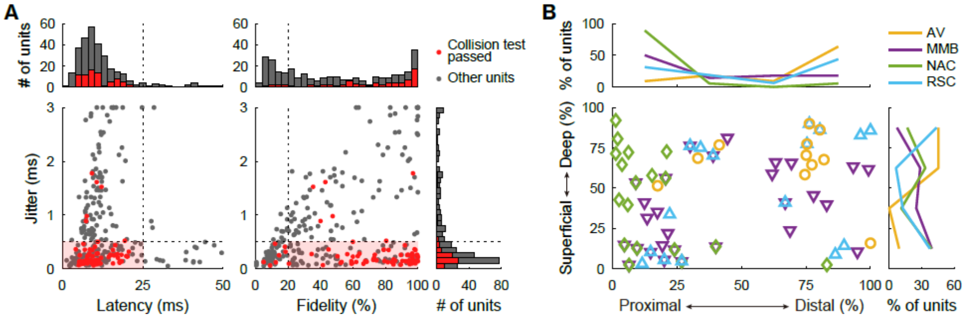


**Fig. S7. Optogenetic identification of projection targets**

(A) Distribution of jitter, latency, and fidelity of evoked spikes in response to blue light stimulation. Responses from stimulating all the target areas (*i.e.*, AV, MMB, NAC, and RSC) are included in this figure. Units with low-jitter (<0.5 ms), short-latency (<25 ms), and high-fidelity (>20%) optical responses (shaded areas) were defined as projection neurons. These criteria were determined by the distribution of units that passed the spike-collision test (red dots and bars). Each dot represents a single unit.

(B) Anatomical distribution of identified projection neurons from all rats mapped onto normalized proximal-distal and superficial-deep axes of the subiculum (SUB) cell layer. Among the identified neurons, 3, 7, and 5 neurons were identified to project to two areas: AV and RSC, MMB and NAC, and MMB and RSC, respectively. One neuron was identified to project on three areas: MMB, NAC, and RSC. These double- and triple-projection neurons are shown as overlapping marks. The distribution of identified neurons was largely consistent with the results of retrograde tracing with CTB488 (Figs. 1, S2, S3): AV-projecting neurons were found at the distal-deep cell layer, MMB-projecting neurons at the superficial cell layer, NAC-projecting neurons at the proximal SUB, and RSC-projecting neurons at the distal SUB.

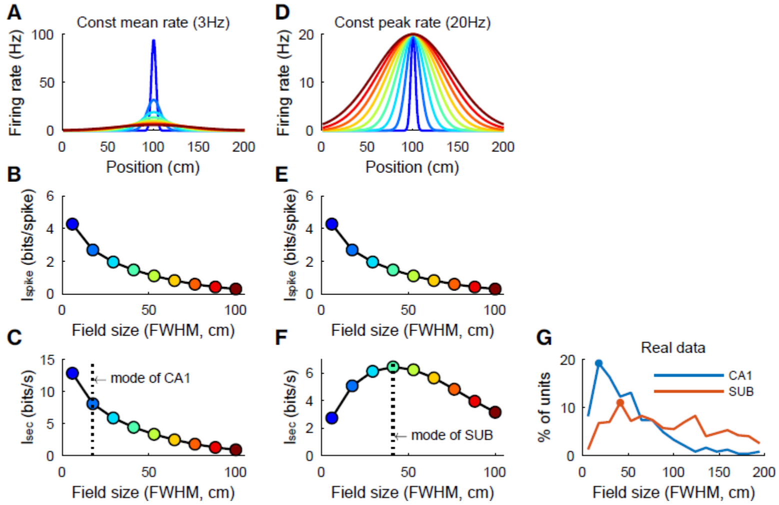


**Fig. S8. Strategy to maximize per-sec spatial information ($I_{sec}$)**

The differences in $I_{spike}$ and mean firing rate and the maintained $I_{sec}$ between the CA1 and SUB (Fig. 2) raises the possibility that these two areas employ different strategies to maximize $I_{sec}$. To clarify the condition that maximizes per-spike spatial information ($I_{spike}$) and $I_{sec}$, we calculated these values for artificial place fields on a linear track (A–F) and compared them with the observed place fields (G). The strategy to maximize $I_{sec}$ with sharp tuning and lower firing rate (A–C) was similar to the observed CA1 representation, while the other strategy to maximize $I_{sec}$ with broader tuning and higher firing rate (D–F) was reminiscent of the observed SUB representation.

(A–C) Gaussian-shaped artificial place fields with variable field sizes [defined as the full-width of half maximum (FWHM) of the tuning curve] were generated under the constraint of constant (3 Hz) mean firing rate (A), and $I_{spike}$ (B) and $I_{sec}$ (C) were calculated according to their definition (see Materials and Methods) with the assumption that time spent in all position bins are uniform. Both $I_{spike}$ and $I_{sec}$ became larger as the field size became smaller.

(D–F) Gaussian-shaped artificial place fields with various field sizes were generated (D) under the constraint of constant (20 Hz) peak firing rate but without the constraint of the mean firing rate. For these place fields, $I_{spike}$ (E) and $I_{sec}$ (F) were calculated. The $I_{spike}$ became larger as the field size became smaller. In contrast, the $I_{sec}$ was largest for the modestly broad, higher mean-firing-rate place field (~41-cm field size for the 200-cm linear track).

(C, F) Dotted lines represent modes of the observed CA1 (C) and SUB (F) field size distribution shown in (G).

(G) Distribution of the observed CA1 and SUB field sizes. Dots on the plots represent modes of distribution.

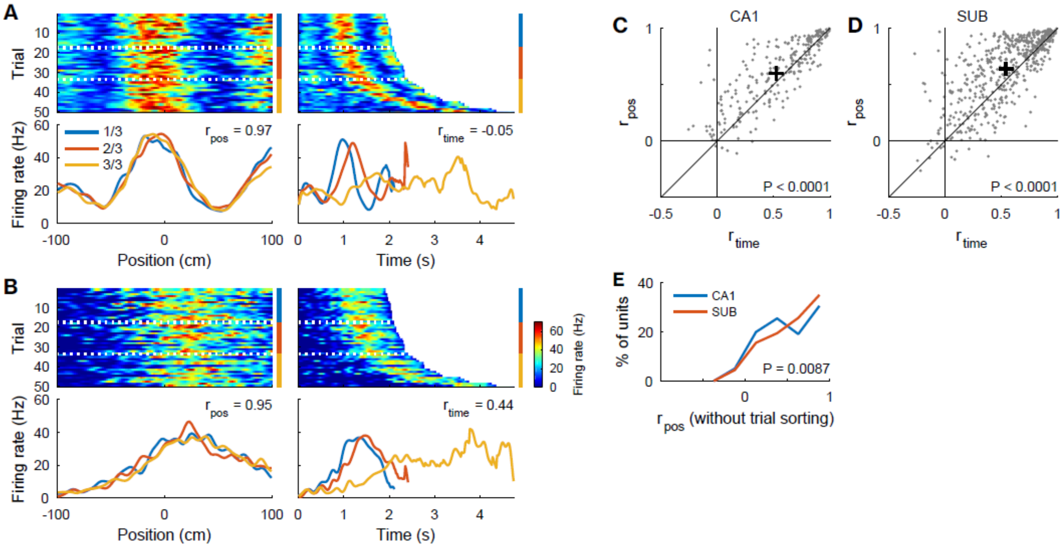


**Fig. S9. Subiculum (SUB) neurons represent place rather than elapsed time on a linear track.**

Hippocampus contains neurons that respond to elapsed time from behaviorally salient timing (*5*). Thus, we investigated which of the positions and elapsed time are better represented by SUB neurons on a linear track.

(A, B) Trial-to-trial rate maps of two representative SUB neurons for eastbound trials (top) plotted according to the animal's position (left) or elapsed time from the start of trials (right). Trials were sorted by their duration (top). Firing rates were averaged over each one-third of the trials (bottom, dotted lines in the top panels). $r_{pos}$ and $r_{time}$ indicate the mean correlation coefficients between all combinations of the three firing rate maps.

(C, D) $r_{pos}$ was larger than $r_{time}$ both in CA1 neurons ($Z = 6.20$, $P < 0.0001$, Wilcoxon signed rank test) (C) and SUB neurons ($Z = 9.45$, $P < 0.0001$) (D), indicating that both CA1 and SUB neurons respond better to animal positions rather than to the elapsed time during running on a linear track. Dots, single directions (eastbound or westbound) from single neurons. Plus signs, means.

(E) Distribution of $r_{pos}$. To estimate the temporal stability of spatial firing, $r_{pos}$ was calculated using the rate maps with the original temporal order without any trial sorting. $r_{pos}$ was larger for SUB neurons than CA1 neurons ($Z = 2.62$, $P = 0.0087$, Wilcoxon rank sum test), suggesting that the spatial tuning in the SUB is more temporally stable than that in the CA1.

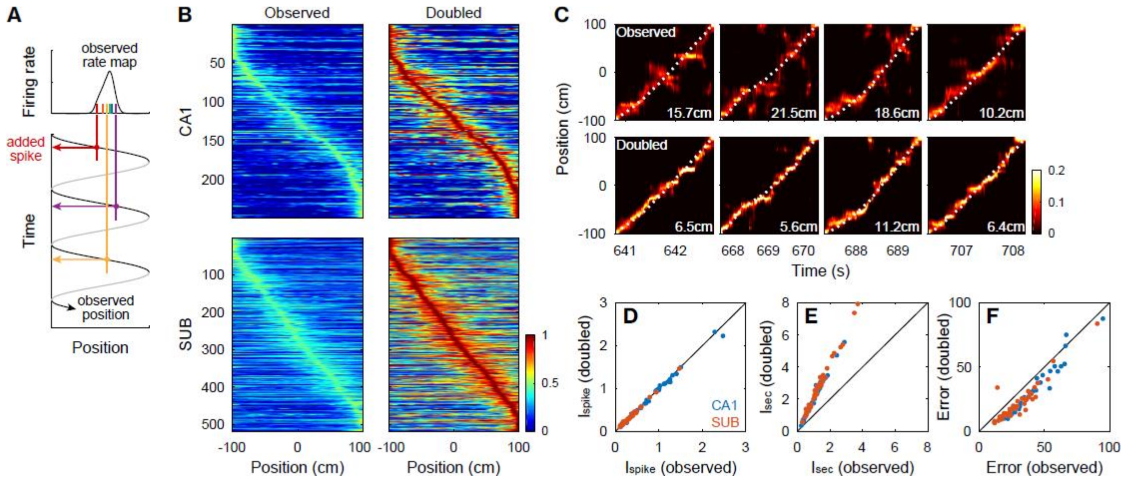


**Fig. S10. Increasing per-sec spatial information ($I_{sec}$) without changing per-spike spatial information ($I_{spike}$) improves position decoding**

We investigated whether increasing $I_{sec}$ by adding spikes improves position decoding performance on a linear track.

(A) Schematic of spike addition to increase $I_{sec}$. Spike addition to each direction (eastbound or westbound) of each neuron without changing spatial tuning was performed in two steps. First, for each direction of a neuron, we generated random spike positions (color raster plot) following the firing rate distribution of the observed rate map (top), so that the generated spikes rarely changed the shape of the rate map. Then, the corresponding spike timing was determined as one of the timing that the rat crossed the generated spike position (bottom). The same number of spikes with the observed spikes were generated for each direction of each neuron and added to the observed spike train, so that the mean firing rate was doubled.

(B) The original observed (left) and spike-doubled (right) firing rate maps of the CA1 (top) and subiculum (SUB; bottom) on a linear track. By spike addition, the firing rates were doubled, while spatial firing was maintained. The color scale represents the firing rate normalized to the peak firing rate of the spike-doubled data. Rate maps were sorted by their peak positions.

(C) Examples of position decoding of four consecutive trials (from left to right) using the observed (top) and spike-doubled (bottom) SUB population activity. Dotted curves: observed animal positions. Numbers: decoding errors.

(D, E) The spike addition did not affect $I_{spike}$ (CA1, 96.9 ± 6.1%; SUB, 100.8 ± 9.3%; mean ± standard deviation; D) but increased $I_{sec}$ (CA1, 197.2 ± 19.5%, $Z$ = 4.94, $P$ < 0.0001; SUB, 208.3 ± 16.4%, $Z$ = 5.65, $P$ < 0.0001; E).

(F) The spike addition decreased decoding error for both CA1 (72.5 ± 15.4%, $Z$ = 4.77, $P$ < 0.0001) and SUB (71.4 ± 29.7%, $Z$ = 5.12, $P$ < 0.0001). These results suggest a tight association between $I_{sec}$ and population decoding performance.

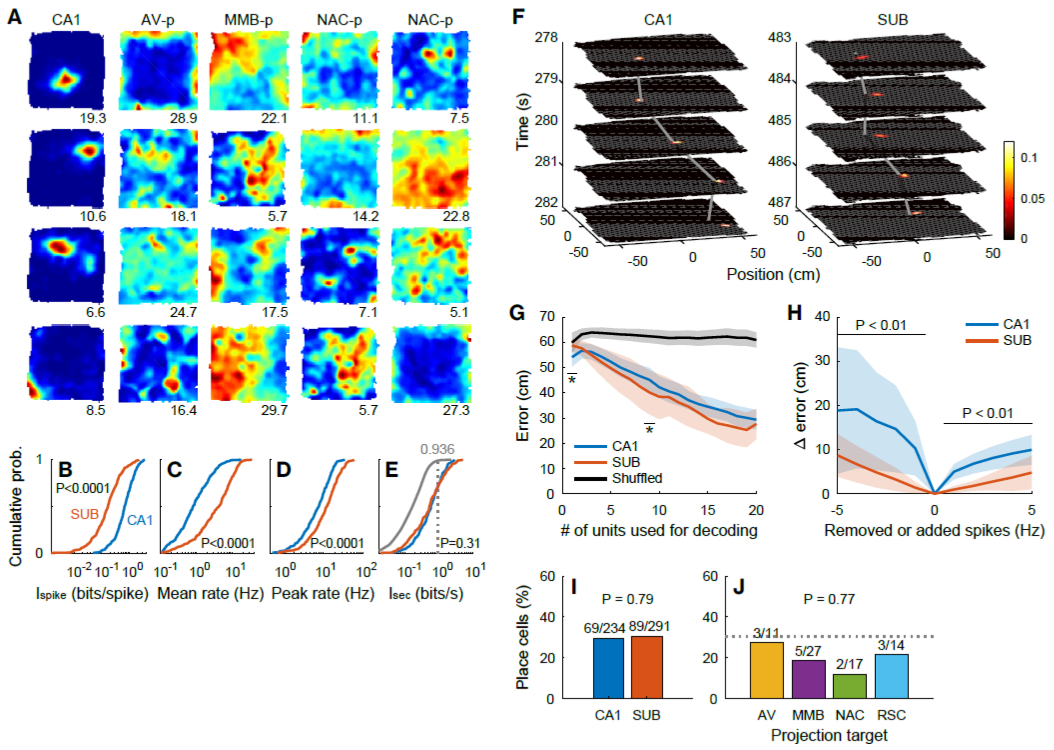

**Fig. S11. Place representation in the CA1 and subiculum (SUB), and identified SUB projection neurons in an open field**

(A) Example rate maps of CA1 (leftmost column) and SUB projection neurons (from the second to the last columns). Numbers indicate the peak firing rate (Hz) of the rate map.

(B–E) Distribution of per-spike spatial information ($I_{spike}$; CA1, 0.89 ± 0.68 bits/spike; SUB, 0.29 ± 0.38 bits/spike; $Z$ = 13.43, $P$ < 0.0001; B), mean firing rate (CA1, 1.5 ± 1.9 Hz; SUB, 5.6 ± 5.2 Hz; $Z$ = 11.48, $P$ < 0.0001; C), peak firing rate (CA1, 9.2 ± 6.4 Hz; SUB, 15.2 ± 11.2 Hz; $Z$ = 6.56, $P$ < 0.0001; D), and per-second spatial information ($I_{sec}$; CA1, 0.78 ± 0.61 bits/s; SUB, 0.82 ± 0.86 bits/s; $Z$ = 1.03, $P$ = 0.31; E) of CA1 (blue) and SUB (red) neurons. P values: Wilcoxon rank sum test. (E) Gray curve shows the distribution of the $I_{sec}$ of shuffled data. Number and dotted line show the 99th percentile of the shuffled data distribution, which was used to define place cells in (I, J).

(F) Examples of position decoding from CA1 (left) or SUB (right) neurons. Color maps: probabilities of animal positions estimated by decoding analysis. Gray lines: observed animal positions.

(G) The decoding error of the SUB (orange) was modestly smaller than that of the CA1 (blue) (main effect of groups, $F_{1,\,445}$ = 25.35, $P$ < 0.0001, two-way ANOVA). Decoding was performed using data during running periods (running speed > 5 cm/s). Black line: chance level of the decoding error estimated by a shuffling procedure. Mean (solid lines) ± standard deviation (SD) (shaded areas). * $P$ < 0.05, CA1 *vs.* SUB, *post-hoc* Bonferroni test.

(H) Differences in decoding errors after randomly removing (negative x values) or adding (positive x values) spikes to each neuron at certain frequencies. The decoding was more resistant to additive noises for SUB neurons than for CA1 neurons. $P$ values, Bonferroni test after two-way repeated-measures analysis of variance. Mean (solid lines) ± SD (shaded areas).

(I, J) Percentage of place cells were similar between the CA1 and SUB neurons ($\chi^2$ = 0.07, $P$ = 0.78) (I), and among groups of SUB projection neurons ($\chi^2$ = 1.13, $P$ = 0.78) (J). Numbers: number of place cells out of the total number of neurons. Dotted line in (J): percentage of place cells in SUB neurons. $P$ values, $\chi^2$ test.

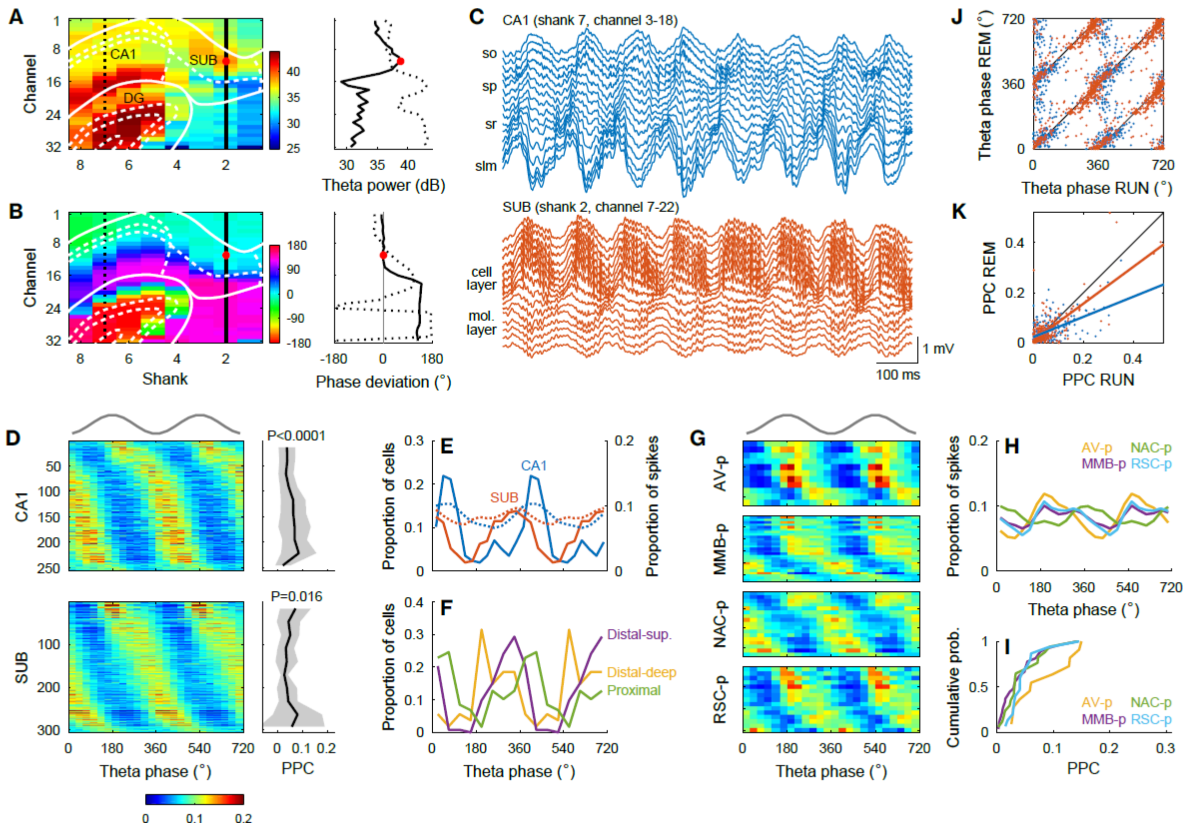


**Fig. S12. Projection-specific theta phase locking during rapid eye movement (REM) sleep**

(A, B) Representative two-dimensional distribution of theta power (A) and theta phase deviation (B) during REM sleep (left). Theta power in the subiculum (SUB) was highest approximately at the center of the SUB cell layer and was comparable with that in the CA1 pyramidal cell layer during both REM sleep (A) and RUN periods (Fig. 5A). Theta phases were almost uniform in the SUB cell layer but shifted ~120 degrees in the SUB molecular layer (B) (Fig. 5B), which was similar with theta phase shifts from the CA1 pyramidal cell layer to the CA1 stratum lacunosum-moleculare (*6*) (B) (Fig. 5B). White lines show the anatomical borders of the CA1, SUB, and dentate gyrus (DG). Values along the CA1 (dotted line) and SUB (solid line) are shown in the right panels. The recording site with the maximal theta power in the SUB cell layer (red dots) was used as reference.

(C) Representative traces of wide-band local field potentials simultaneously recorded from the CA1 (top) and SUB (bottom) during REM sleep. so, stratum oriens; sp, stratum pyramidale; sr, stratum radiatum; slm, stratum lacunosum-moleculare.

(D) Spike phase distribution along reference SUB theta oscillations (left) and pairwise phase consistency (PPC) (right) in the CA1 (top) and SUB (bottom) during REM sleep. Most neurons in the CA1 (89.2%) and SUB (97.8%) were significantly phase-locked ($P < 0.05$, Rayleigh test) to reference SUB theta oscillations, and only these neurons are shown in this figure. P values, one-way analysis of variance. Mean (solid line) ± standard deviation (SD; shaded areas). Considering the entire population, PPC was modestly smaller in SUB neurons than in CA1 neurons (CA1, 0.054 ± 0.050; SUB, 0.049 ± 0.064; Z = 2.57; $P = 0.010$). (D, G) The top gray trace indicates idealized reference theta cycles in the SUB cell layer.

(E) Distribution of preferred theta phases (left y axis, solid lines) and spike theta phases (right y axis, dotted lines) of CA1 (blue) and SUB (orange) neurons. The preferred theta phases for SUB neurons were earlier than those for CA1 neurons (CA1, 52.5 ± 60.2 degrees; SUB, 324.6 ± 64.5 degrees; $U^2$ = 2.30, $P$ < 0.001, Watson $U^2$ test).
(F) Distribution of preferred theta phases for SUB neurons located at the proximal (green; proximal one-third of the cell layer; 24.6 ± 64.5 degrees), distal-deep (yellow; distal two-thirds and deep one-third of the cell layer; 257.1 ± 60.8 degrees), and distal-superficial (purple; distal two-thirds and superficial two-thirds of the cell layer; 313.9 ± 49.7 degrees) part of the SUB cell layer. These three distributions were significantly different with each other ($P$ < 0.01, Watson $U^2$ test with Bonferroni correction).
(G) Spike phase distribution of SUB projection neurons along the reference SUB theta oscillations.
(H, I) Distribution of spike theta phases (H) and PPC (I) of the SUB projection neurons during RUN periods. The preferred theta phases of NAC-p neurons were different from those of AV-p neurons ($P$ = 0.042, Watson $U^2$ test with Bonferroni correction). Circular means ± angular deviations of preferred theta phases were as follows: AV-p, 234.3 ± 51.1 degrees; MMB-p, 251.8 ± 60.1 degrees, NAC-p, 12.8 ± 61.4 degrees, and RSC-p, 267.9 ± 55.0 degrees.
(J, K) Relationship of preferred theta phases (J) and PPC (K) during RUN periods and REM sleep. Dots represent single neurons in the CA1 (blue) and SUB (orange). Black lines show diagonal. (J) Consistent with a previous report (*43*), a fraction of CA1 neurons (16.9%) shifted their preferred theta phases by 90–270 degrees during REM sleep compared with the RUN periods (Fig. 5J). In contrast, a significantly smaller proportion of SUB neurons (6.6%) showed REM shifting ($\chi^2$ = 13.74, $P$ = 0.0002). The preferred theta phases of SUB neurons were correlated between RUN and REM periods (circular-circular correlation = 0.69, $P$ < 0.0001). (K) PPC during RUN and REM periods were correlated in both CA1 ($r$ = 0.54, $P$ < 0.0001) and SUB ($r$ = 0.77, $P$ < 0.0001) neurons. Color lines: linear regression.

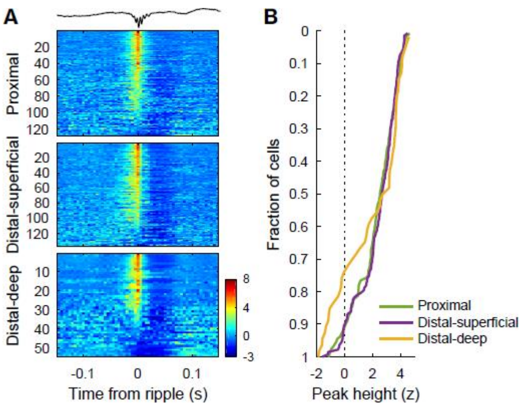


**Fig. S13. Location-specific firing modulation by sharp-wave/ripples (SPW-Rs) during slow-wave sleep.**

(A) Ripple-triggered average of firing rates of individual subiculum (SUB) neurons located at the proximal (top; proximal one-third of the cell layer), distal-superficial (middle; distal two-thirds and superficial two-thirds of the cell layer), and distal-deep (bottom; distal two-thirds and deep one-third of the cell layer) part of the SUB cell layer. Color maps represent z-scored firing rates. Time zero indicates the negative peak of ripple-band (140–230 Hz) local field potential traces of the reference recording site in the SUB. Topmost trace indicates an example SPW-Rs trace in the SUB.

(B) Mean z-scored firing rate averaged around the negative ripple peaks (–10 to 10 ms). Most SUB neurons were significantly ($P < 0.01$, $t$-test) activated during ripples (proximal: 75.0%, distal-superficial: 77.4%, distal-deep: 61.1%). Neurons that were significantly ($P < 0.01$, $t$-test) suppressed during ripples were concentrated in the distal-deep part of the SUB cell layer ($P < 0.001$ by $\chi^2$ test, proximal: 2.3%, distal-superficial: 1.5%, distal-deep: 16.7%).

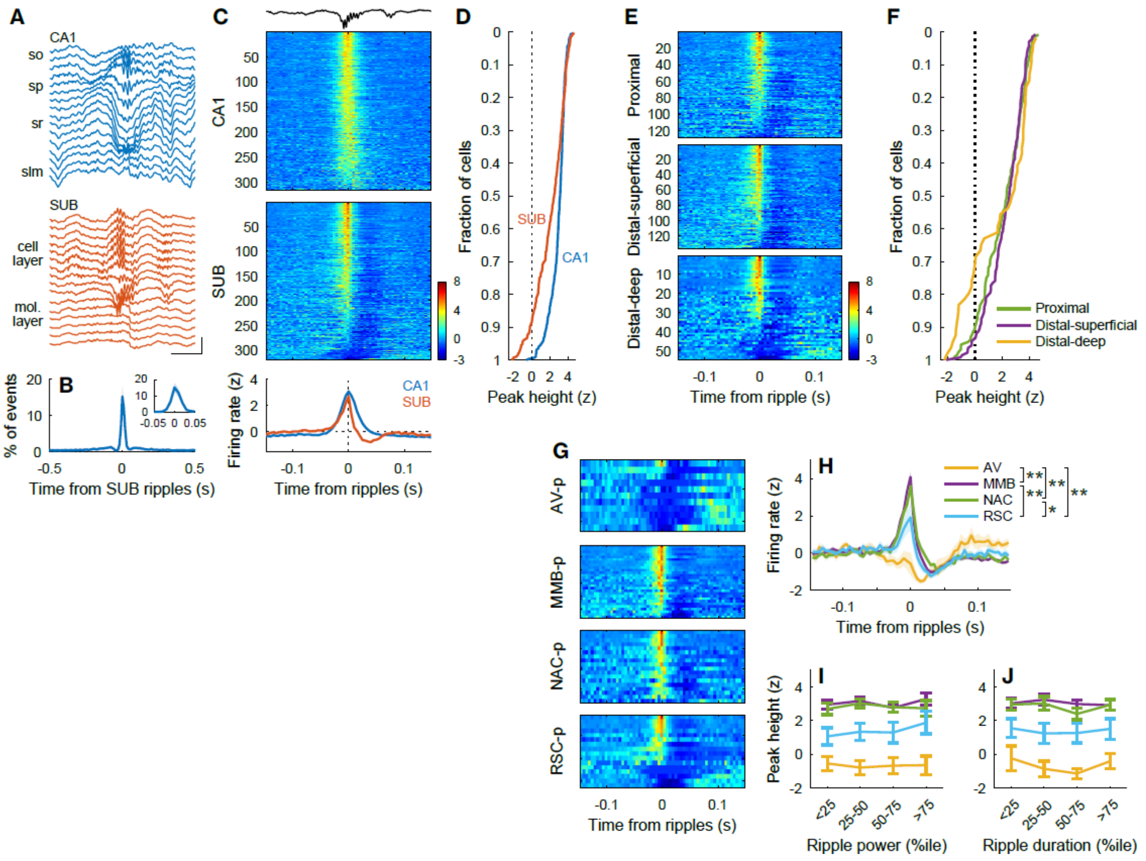

**Fig. S14. Projection-specific firing modulation by sharp-wave/ripples (SPW-Rs) during awake rest periods.**

(A) Representative SPW-Rs during awake rest (REST) periods in the CA1 (top) and subiculum (SUB; bottom). Simultaneously recorded, wide-band local field potential (LFP) traces are shown. Scale bar: 50 ms, 1 mV. so, stratum oriens; sp, stratum pyramidale; sr, stratum radiatum; slm, stratum lacunosum-moleculare.

(B) Cross-correlogram (CCG) of ripple events detected in the CA1 pyramidal cell layer and SUB cell layer. Inset shows the same CCG at a finer temporal scale. Mean (solid line) ± standard deviation (shaded area).

(C) Ripple-triggered average of firing rates of individual CA1 (top) and SUB (middle) neurons, and the firing rate traces averaged over CA1 or SUB neurons (bottom). Color maps represent z-scored firing rates. Time zero indicates the negative peak of ripple-band (140–230 Hz) LFP traces of the reference recording site in the SUB. Topmost trace indicates an example SPW-Rs trace in the SUB. Ripple events that were detected in single recording sites at the center of the SUB cell layer were used as a reference.

(D) Mean z-scored firing rate averaged around negative ripple peaks (–10 to 10 ms). Some SUB neurons (6.0%) were significantly suppressed during ripples, while no CA1 neurons (0%) were suppressed ($\chi^2$ = 19.31, $P$ < 0.0001, $\chi^2$ test).

(E) Ripple-triggered average of firing rates of individual SUB neurons located at the proximal (top), distal-superficial (middle), and distal-deep (bottom) part of the SUB cell layer. Color maps represent z-scored firing rates.

(F) Mean z-scored firing rate averaged around negative ripple peaks (–10 to 10 ms). Most SUB neurons were significantly ($P$ < 0.01, $t$-test) activated during ripples (proximal: 64.8%, distal-superficial: 73.0%, distal-deep: 55.6%). Neurons that were significantly ($P$ < 0.01, $t$-test) suppressed during ripples were concentrated in the distal-deep part of the SUB cell layer ($P$ < 0.001 by $\chi^2$ test, proximal: 5.5%, distal-superficial: 1.5%, distal-deep: 18.5%).
(G) Color maps of ripple-triggered average of z-scored firing rates of identified SUB projection neurons.
(H) Ripple-triggered average of z-scored firing rates of SUB projection neurons. * $P$ < 0.05, ** $P$ < 0.01, Tukey test for peak height. Mean (solid lines) ± standard error of the mean (SEM) (shaded areas).
(I, J) Peak height of firing rate (z-scored) as a function of ripple power (I) and ripple duration (J). The plot colors are the same as in (H). (I) Two-way repeated-measures analysis of variance (ANOVA) revealed the main effect of projection neuron groups ($F_{3, 69}$ = 17.96, $P$ < 0.001). $P$ < 0.01 by *post-hoc* Bonferroni test, AV-p *vs.* all other projection neuron groups, MMB-p *vs.* RSC-p. (J) Two-way repeated-measures ANOVA revealed the main effect of projection neuron groups ($F_{3, 69}$ = 18.03, $P$ < 0.001). $P$ < 0.01 by *post-hoc* Bonferroni test, AV-p *vs.* all other projection neuron groups, MMB-p *vs.* RSC-p. $P$ < 0.05, NAC-p *vs.* RSC-p. Mean ± SEM.
